# Comparison of R9.4.1/Kit10 and R10/Kit12 Oxford Nanopore flowcells and chemistries in bacterial genome reconstruction

**DOI:** 10.1101/2022.04.29.490057

**Authors:** Nicholas Sanderson, Natalia Kapel, Gillian Rodger, Hermione Webster, Samuel Lipworth, Teresa street, Tim Peto, Derrick Crook, Nicole Stoesser

**Author notes:** Corresponding/alternative corresponding author Corresponding authors Nicholas Sanderson Nicole Stoesser. Contributed equally (Nicholas Sanderson, Natalia Kapel). **Repositories:** Nanopore fast5 and fastq data are available in the ENA under project accession: PRJEB51164. Assemblies have been made available at: https://figshare.com/articles/online_resource/q20_comparison_genome_assemblies/19683867 Code and analysis outputs are available at: https://gitlab.com/ModernisingMedicalMicrobiology/assembly_comparison_analysis/-/tree/main (tagged version v0.5.5).

## Abstract

2.

Complete, accurate, cost-effective, and high-throughput reconstruction of bacterial genomes for large-scale genomic epidemiological studies is currently only possible with hybrid assembly, combining long- (typically using nanopore sequencing) and short-read (Illumina) datasets. Being able to utilise nanopore-only data would be a significant advance. Oxford Nanopore Technologies (ONT) have recently released a new flowcell (R10.4) and chemistry (Kit12), which reportedly generate per-read accuracies rivalling those of Illumina data. To evaluate this, we sequenced DNA extracts from four commonly studied bacterial pathogens, namely *Escherichia coli*, *Klebsiella pneumoniae*, *Pseudomonas aeruginosa* and *Staphylococcus aureus*, using Illumina and ONT’s R9.4.1/Kit10, R10.3/Kit12, R10.4/Kit12 flowcells/chemistries. We compared raw read accuracy and assembly accuracy for each modality, considering the impact of different nanopore basecalling models, commonly used assemblers, sequencing depth, and the use of duplex versus simplex reads. “Super accuracy” (sup) basecalled R10.4 reads - in particular duplex reads - have high per-read accuracies and could be used to robustly reconstruct bacterial genomes without the use of Illumina data. However, the per-run yield of duplex reads generated in our hands with standard sequencing protocols was low (typically <10%), with substantial implications for cost and throughput if relying on nanopore data only to enable bacterial genome reconstruction. In addition, recovery of small plasmids with the best-performing long-read assembler (Flye) was inconsistent. R10.4/Kit12 combined with sup basecalling holds promise as a singular sequencing technology in the reconstruction of commonly studied bacterial genomes, but hybrid assembly (Illumina+R9.4.1 hac) currently remains the highest throughput, most robust, and cost-effective approach to fully reconstruct these bacterial genomes.

**Impact statement:** Our understanding of microbes has been greatly enhanced by the capacity to evaluate their genetic make-up using a technology known as whole genome sequencing. Sequencers represent microbial genomes as stretches of shorter sequence known as ‘reads’, which are then assembled using computational algorithms. Different types of sequencing approach have advantages and disadvantages with respect to the accuracy and length of the reads they generate; this in turn affects how reliably genomes can be assembled.

Currently, to completely reconstruct bacterial genomes in a high-throughput and cost-effective manner, researchers tend to use two different types of sequencing data, namely Illumina (short-read) and nanopore (long-read) data. Illumina data are highly accurate; nanopore data are much longer, and this combination facilitates accurate and complete bacterial genomes in a so-called “hybrid assembly”. However, new developments in nanopore sequencing have reportedly greatly improved the accuracy of nanopore data, hinting at the possibility of requiring only a single sequencing approach for bacterial genomics.

Here we evaluate these improvements in nanopore sequencing in the reconstruction of four bacterial reference strains, where the true sequence is already known. We show that although these improvements are extremely promising, for high-throughput, low-cost complete reconstruction of bacterial genomes hybrid assembly currently remains the optimal approach.

**Data summary:** The authors confirm all supporting data, code and protocols have been provided within the article, through supplementary data files, or in publicly accessible repositories.

Nanopore fast5 and fastq data are available in the ENA under project accession: PRJEB51164.

Assemblies have been made available at: https://figshare.com/articles/online_resource/q20_comparison_genome_assemblies/196838 67.

Code and analysis outputs are available at: https://gitlab.com/ModernisingMedicalMicrobiology/assembly_comparison_analysis/-/tree/main (tagged version v0.5.5).

## 5. Introduction

Bacterial whole genome sequencing has become a prominent tool in the biological sciences, with wide-ranging applications from epidemiology to diagnostics(1). Important considerations include sequencing throughput, read length (which facilitates complete reconstruction of bacterial chromosomes and plasmids), read accuracy, accessibility and cost. Historically, short-read Illumina sequencing has been the leading high-throughput, high-accuracy technology, but is limited in its capacity to completely reconstruct genomes, particularly in the presence of repetitive sequences. Nanopore sequencing (Oxford Nanopore Technologies [ONT]) has become one of the most widely adopted long-read sequencing approaches, enabled by affordable, small-footprint sequencing platforms, but has been limited to some extent by its accuracy. Combining short and long-read sets from both technologies in the form of hybrid assembly has facilitated cost-effective, highly accurate and scalable genome reconstruction for large bacterial isolate collections(2, 3), such as by multiplexing 96 *E. coli* isolates on a single nanopore flowcell(3). For nanopore sequencing, developments in multiplexing, rapid library preparation and flow cell reuse after washing have streamlined this process(4).

ONT have undertaken iterative development of their sequencing flowcells and chemistries, releasing the R10.3 (FLO-MIN111) flowcells for consumers in January 2020 and the Kit12 (Q20+) chemistry and R10.4 flowcell (FLO-MIN112) in their store in late 2021. The proposed advantages of the R10.4/Kit12 system include: (i) a new motor to facilitate more controlled passage of the nucleic acid template through the sequencing pore thereby avoiding template slippage; (ii) “duplex” read sequencing - where the forward and reverse strand of a single nucleic acid molecule are sequenced in succession to improve accuracy; and (iii) an optimized pore with a longer pore head to better resolve homopolymers.

These new developments however come with some potential disadvantages. Sequencing yields for the R10.3 flowcells were lower than those using R.9.4.1 flowcells (thought to be due to the slower passage of template through pores)(5). The use of R10 flowcells also currently requires a ligation-based library preparation, which results in longer sequencing turnaround times when compared with rapid transposase-based library preparation kits which can be used with R9.4.1 flowcells. Ligation-based preparations may also miss the capture and sequencing of small plasmids(6). The reported improvements in per-read accuracy with R10/Kit12 are also potentially dependent on the use of super accuracy (sup) basecalling models; however, on the same computing infrastructure sup basecalling takes 2-8x longer than the previous typical approach using high accuracy (hac) basecalling models, which may preclude “on-machine” basecalling in real-time(7).

Sequencing accuracy can be characterized using several different metrics, including: (i) raw read accuracy (the accuracy achieved when reading a single nucleic acid fragment once) and (ii) assembly accuracy (the capacity to accurately reconstruct complete genomes in terms of structure, sequence identity and coding sequence content). We therefore set out to compare data and assemblies generated by R9.4.1/Kit10 and R10/Kit12 nanopore flowcells/chemistries, comparing these with Illumina-only sequence data and hybrid assembly, and investigating the impact of sup versus hac basecalling and metrics for duplex sequencing reads. We undertook this comparison for four reference bacterial strains reflecting different species, genome sizes, %GC content, plasmid content and plasmid sizes. We also evaluated the impact of sequencing depth on the capacity to reconstruct the reference bacterial genomes, and whether flowcell washing would still enable flow cell reuse with the new flowcells and chemistry.

## 6. Methods

### Bacterial isolates and DNA extraction

Four reference bacterial strains were sequenced for this study, namely: *Escherichia coli* CFT073 (Genbank accession: NC_004431.1), *Klebsiella pneumoniae* MGH78578 (NC_009648.1-NC_009653.1), *Pseudomonas aeruginosa* PAO1 (NC_002516.2) and *Staphylococcus aureus* MRSA252 (NC_002952.2). Stock cultures were stored at -80°C in nutrient broth supplemented with 10% glycerol. For DNA extraction, stocks were sub-cultured on Columbia blood agar at 37°C overnight.

Long fragment DNA extraction from sub-cultured strains was performed using the Qiagen Genomic tip 100/G kit (Qiagen). Quality and fragment length assessments were measured with the Qubit fluorometer (ThermoFisher Scientific) and TapeStation (Agilent). The same DNA extract, stored in elution buffer at 4°C was used for all sequencing experiments. DNA concentration and fragment lengths were evaluated longitudinally to ensure that there was minimal obvious degradation (Tables S1-4, Figs.S1-3).

### Nanopore sequencing

The experimental workflow is shown in Fig.1. For the experiment using the R9.4.1 (FLO-MIN106) flowcell (denoted as R.9.4 throughout), ONT sequencing libraries were prepared by multiplexing DNA extracts from all four isolates using the Rapid Barcoding Sequencing (SQK-RBK004) kit according to the manufacturer’s protocol; sequencing was undertaken on a GridION for 48 hours.

**Figure 1.**
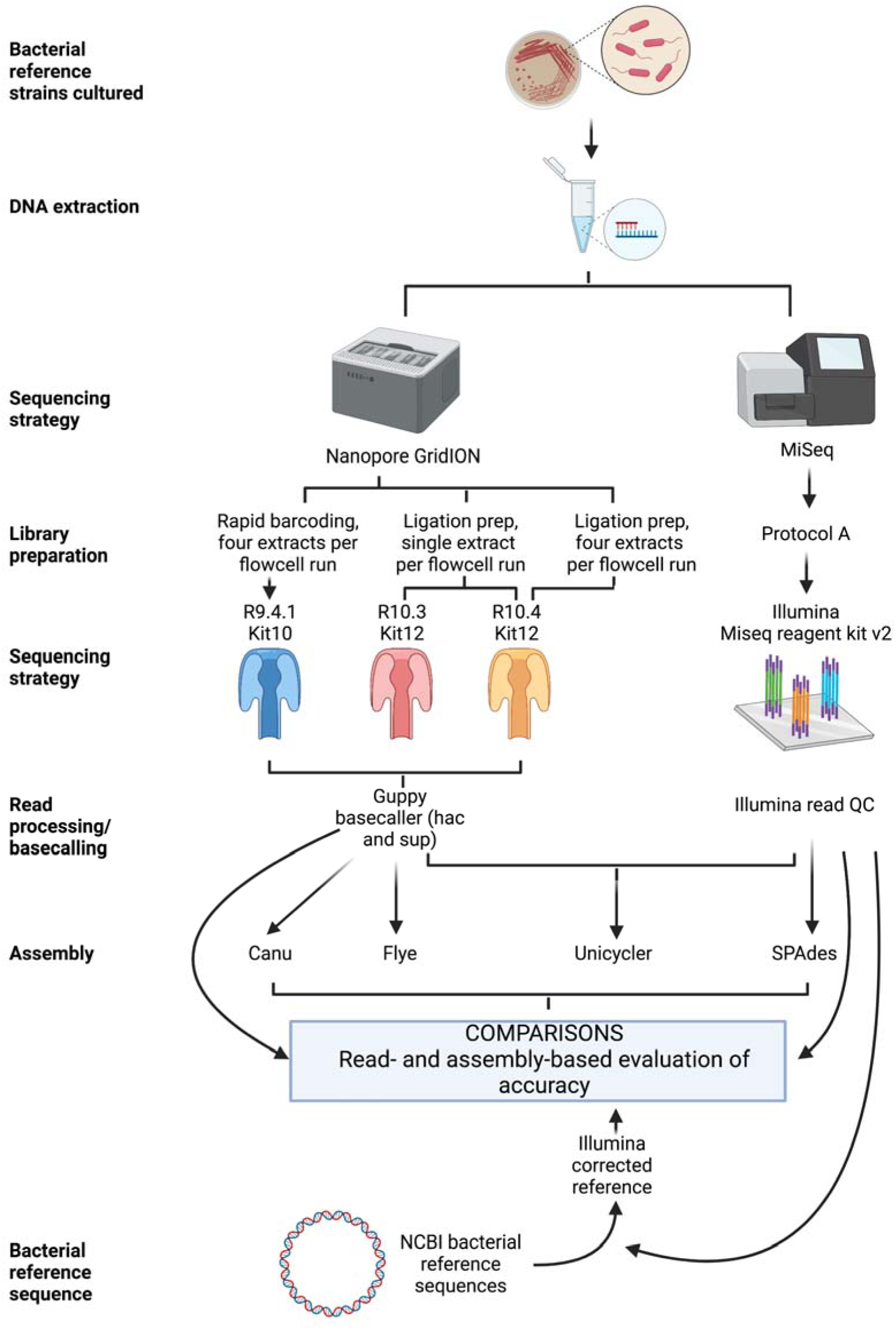
Experimental workflow.

For the experiments using the R10.3 (FLO-MIN111) and R10.4 (FLO-MIN112) flowcells, ONT sequencing libraries were prepared from DNA extracts using the Q20+ Early Access Kit (SQK-Q20EA) ligation-based protocol. During adapter ligation and clean-up the long fragment buffer was used to enrich for DNA fragments >3kb. Each DNA extract was sequenced on a single flowcell. After sequencing the *S. aureus* MRSA252 library, the R10.4 (FLO-MIN112) flowcell was washed with the flowcell wash kit (EXP-WSH004) according to the manufacturer’s protocol, before reusing the flowcell to sequence the *P. aeruginosa* PAO1 library. For the R10.3 experiments, sequencing was undertaken on a GridION for 48 hours; for the unplexed R10.4 experiments sequencing times were terminated prematurely.

The flowcell usage strategy and pore counts for each flowcell prior to use are summarised in Table S5.

Finally, in a separate experiment, the four DNA extracts were also multiplexed on the R10.4 (FLO-MIN112) flowcell using the Native Barcoding Kit (SQK-NBD112.24); sequencing was undertaken on a GridION for 48 hours.

### Illumina sequencing

DNA extracts for all isolates were also sequenced on the Illumina MiSeq, as part of a wider run plexing 20 bacterial extracts. Libraries were constructed following the Illumina DNA Prep protocol, according to the manufacturer’s instructions (including standard normalization for libraries [“Protocol A”]). Library DNA concentrations were quantified by Qubit fluorometry and size distributions of libraries determined using the TapeStation, as above. Sequencing was performed using the MiSeq Reagent Micro Kit v2, generating 150 bp paired-end reads.

### Data processing and bioinformatic methods

R10.4 duplex read pairs were identified and prepared for basecalling using ONT’s duplex tools (https://pypi.org/project/duplex-tools/; v 0.2.9). R9.4, R10.3, and R10.4 raw nanopore reads were hac basecalled with Guppy (ONT) versions 5.0.12+eb1a981 (dna_r9.4.1_450bps_hac.cfg), 5.0.13+bbad529 (res_dna_r103_q20ea_crf_v034.cfg), and 5.0.16+b9fcd7b (dna_r10.4_e8.1_hac.cfg) respectively, as recommended by ONT. R9.4, R10.3, R10.4 (all reads) and R10.4 duplex raw nanopore reads were also basecalled using sup models dna_r9.4.1_e8.1_sup.cfg, dna_r10.3_450bps_sup.cfg, dna_r10.4_e8.1_sup.cfg. Basecalled read summary statistics were generated with seqkit stats using ‘-T’ and ‘-all’ flags(8).

Nanopore reads were subsampled using Rasusa(9) to depths of 10, 20, 30, 40, 50, and 100 average coverage. Nanopore reads were assembled with Canu (version 2.2, using maxInputCoverage=100 and otherwise default parameters)(10), or Flye (using the --meta and --nano-hq parameters and otherwise defaults, version 2.9-b1768)(11), both of which are commonly used long-read only assemblers that have been shown to optimize long-read only assembly quality(12). We also explored the impact of polishing nanopore assemblies with 1, 2 and 3 rounds of Medaka (default settings; https://github.com/nanoporetech/medaka).

Subsampled nanopore reads were combined with Illumina reads for hybrid assembly using Unicycler (version 0.4.8, default parameters)(13). The SPAdes (version 3.15.3)(14) assemblies generated from Illumina data as part of the Unicycler pipeline were used as the Illumina-only assemblies for comparative evaluations.

Given the previous discrepancies observed between multiple resequenced assemblies for *E. coli* CFT073 and *K. pneumoniae* MGH78578(15), and the genetic and phenotypic differences observed in different laboratory sub-culture stocks of *P. aeruginosa* PAO1(16, 17), we generated an Illumina-corrected reference sequence to use as the “gold standard” comparator for this evaluation. Reference genomes for *E. coli* CFT073 (Genbank accession: AE014075.1), *K. pneumoniae* MGH78578 (CP000647.1), *P. aeruginosa* PAO1 (NC_002516.2), *S. aureus* MRSA252 (NC_002952.2) and the respective Illumina datasets generated for this study were used as inputs for the SNIPPY pipeline (version 4.6.0) (https://github.com/tseemann/snippy); output consensus fasta files represented the new Illumina-corrected reference sequences used in this study.

Assembled contigs from nanopore, Illumina, and hybrid assemblies were compared against the Illumina-corrected reference sequences using DNAdiff version 1.3(18).

Assembled contigs from nanopore, Illumina, and hybrid assemblies as well as the Illumina- corrected reference sequences were annotated with Prokka (version 1.14.6)(19), using the corresponding reference GenBank files to ascertain reference proteins using the ‘--proteins’ flag.

Translated amino acid sequences for Prokka annotations in the different test assemblies (Canu, Flye [long-read only], Unicycler [hybrid long-/short-read], SPAdes [short-read only]) and Illumina-corrected reference sequences were compared using the script AAcompare.py in the workflow provided (see below for the repository link). This looked for exact amino acid sequence matches (i.e. 100% identity along 100% of the translated protein) between the Illumina-corrected reference and assembled contigs to determine how intact assembled coding sequences were for each assembly method.

Per read error rates were calculated by mapping the raw reads to the Illumina corrected references sequences using minimap2 (version 2.22-r1101)(18). The percent identity was calculated from the query distance (NM tag) divided by the query length, multiplied by 100, using the bamreadstats.py script provided in the gitlab repository (link below).

A workflow for this analysis has been written using nextflow(18) and is available on gitlab (https://gitlab.com/ModernisingMedicalMicrobiology/assembly_comparison). Outputs from the analyses are also available in this repository (tagged version v0.5.5).

### Data visualization

Figures and plots for this manuscript were generated using the ggplot2 and patchwork packages in R (v3.6.2), and Biorender (www.biorender.com).

## 7. Results

### Sequencing yield and read length distributions

The total data yield after 48 hours of sequencing from the R9.4 flowcell was 11.0Gb (four isolate extracts multiplexed on one sequencing run), compared with 4.0Gb for the R10.4 multiplexed run (Table 1, Fig.S4). For the individual R10.3 flowcells a median of 8.2Gb/flowcell (IQR: 7.3-8.8Gb) were generated by 48 hours of sequencing, and 6.7Gb/flowcell (IQR: 6.6-7.4Gb) for the R10.4 flowcells respectively by 20-30 hours of sequencing (Table 1, Fig.S4). 21.3Mb of data were generated for the extracts from the Illumina runs (Table 1).

**Table 1.**
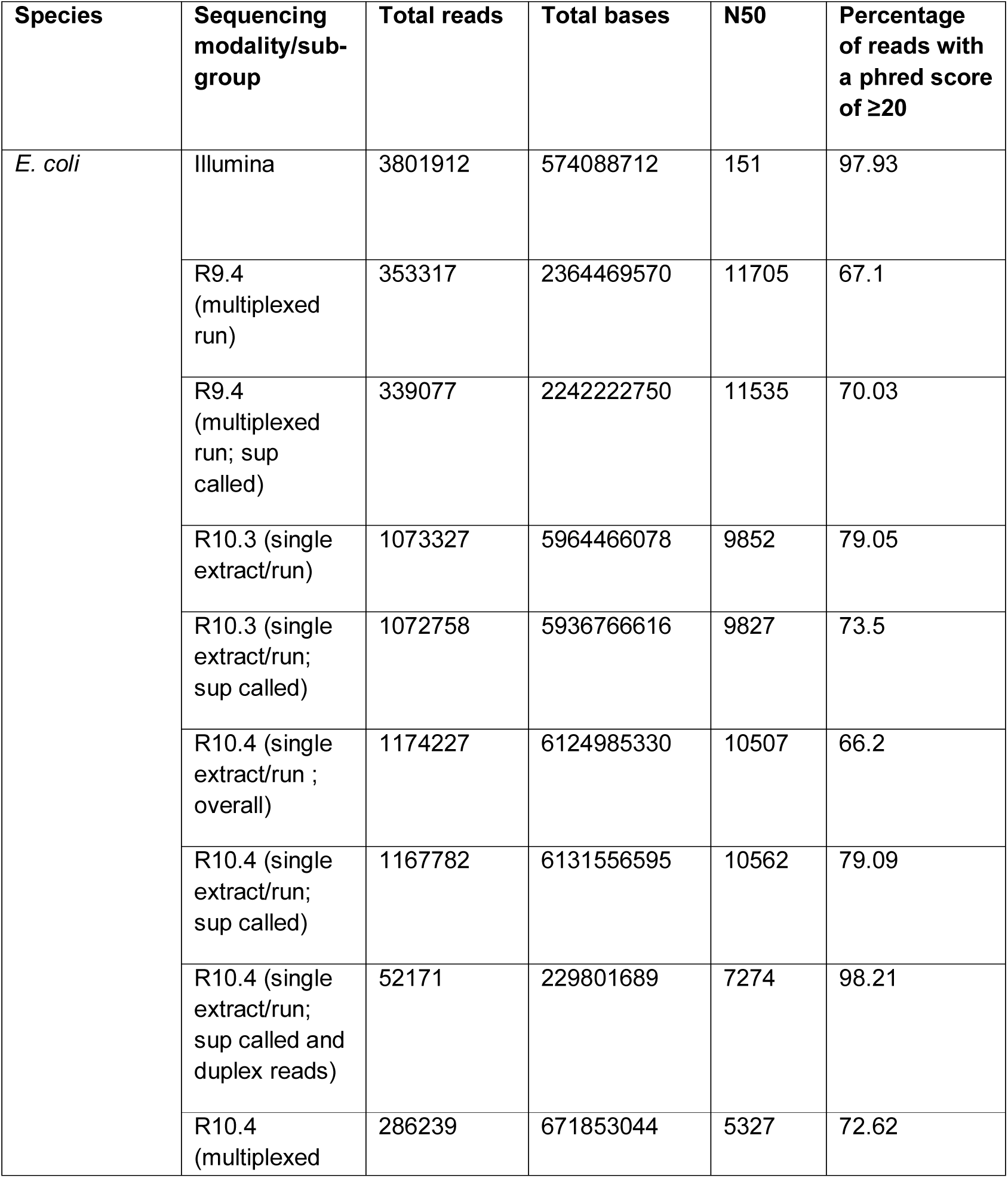
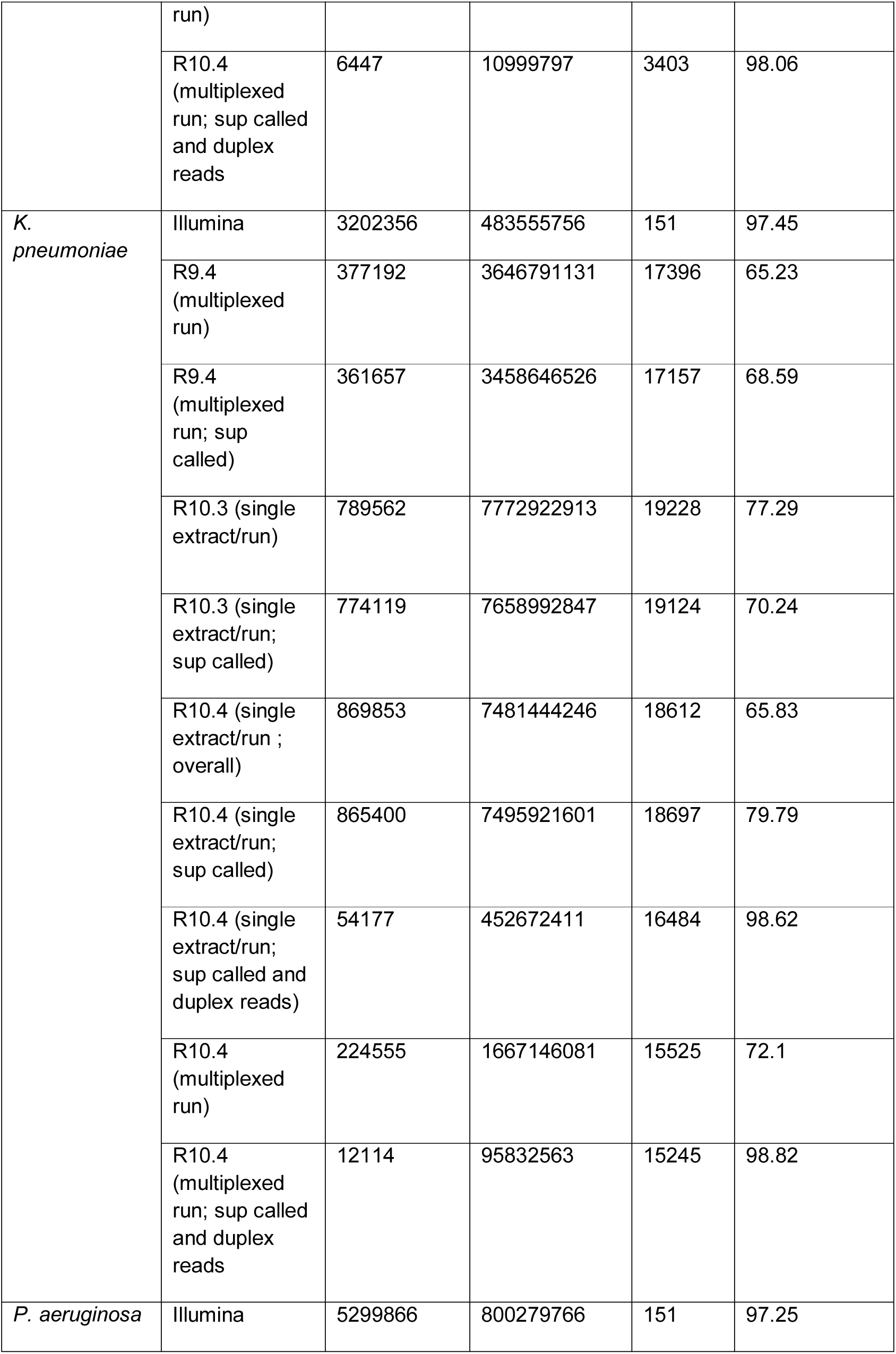
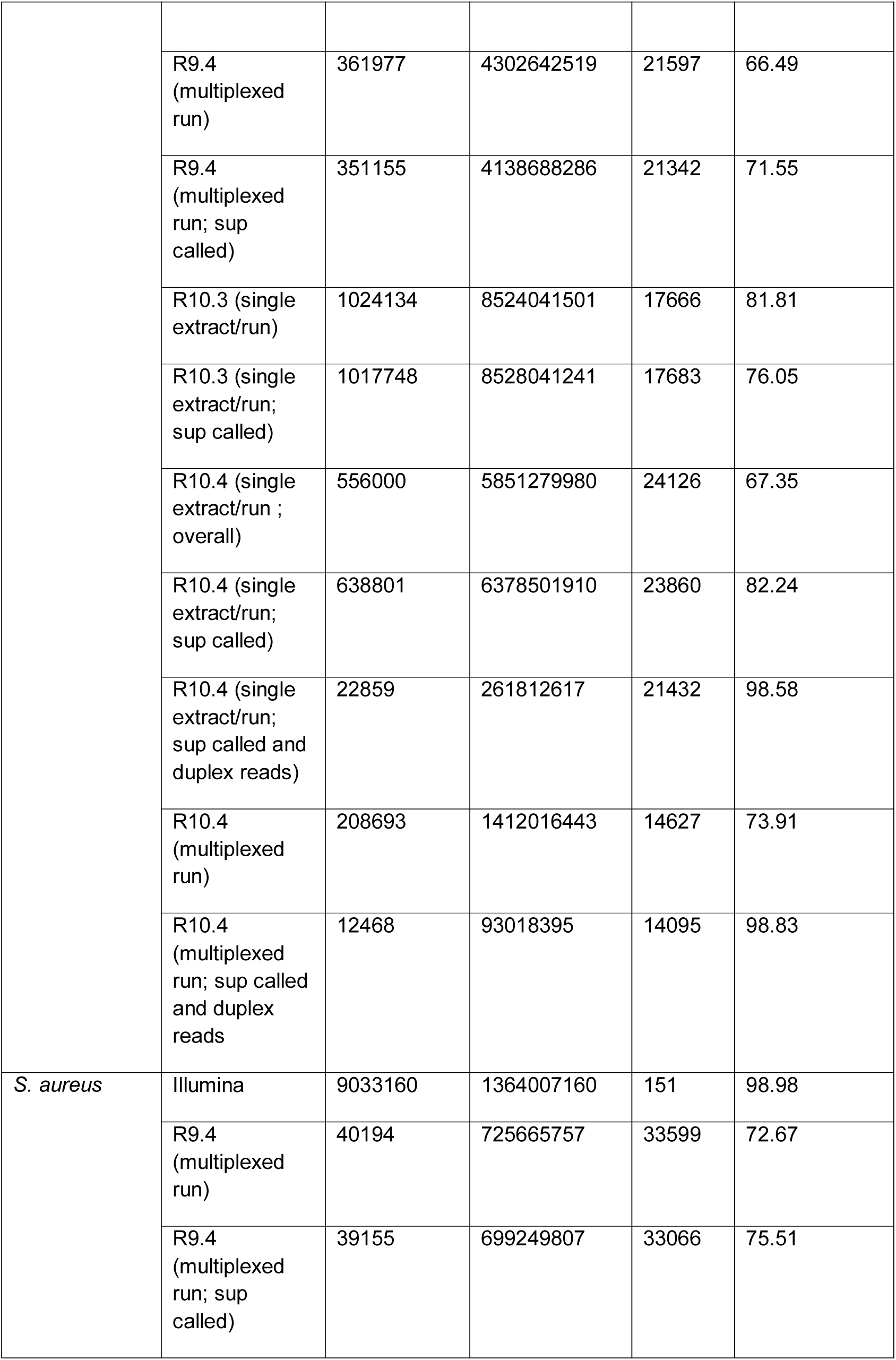
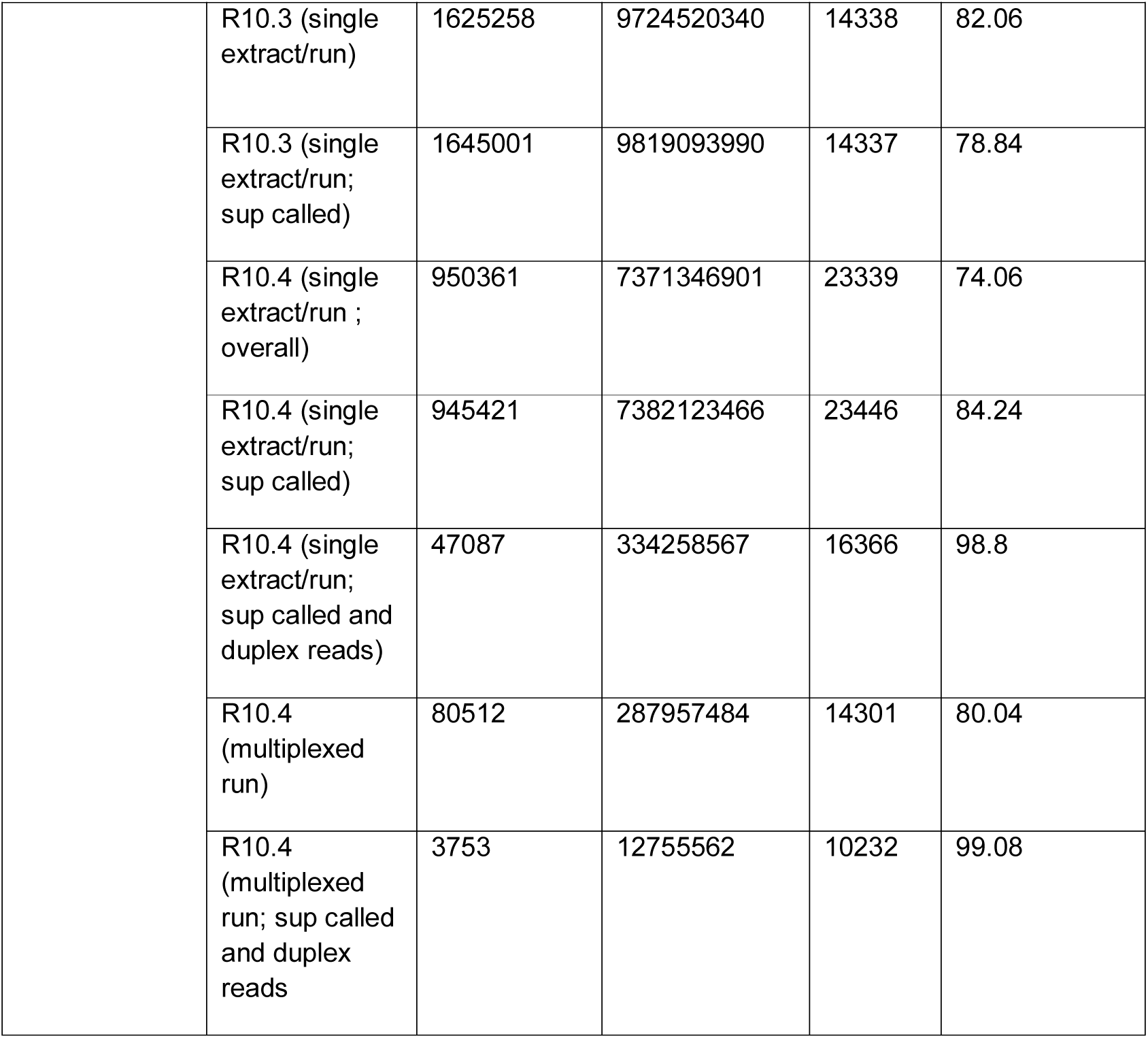
Sequencing read statistics by sequencing modality and bacterial species. Note for R.9.4/Kit10 four isolates were plexed and the total data output is a composite of the individual outputs; for the R10.3/Kit12 and R10.4/Kit12 evaluations each isolate extract was initially run separately. The same flowcell was washed and then re-used for the R10.4 evaluation for the *S. aureus* and then *P. aeruginosa* isolates. Finally, the four DNA extracts were also multiplexed on a single R10.4/Kit12 run.

Read length distributions for a subsample of 1000 reads by modality and species are shown in Fig.2; overall, across species for nanopore data the median read length was 3580bp, the maximum read length 388620bp and the minimum read length 77bp. Median read lengths generated using R9.4 were longer (6273bp versus 2930bp for R10.4; two-sample Wilcoxon test, p<0.001, comparison for hac basecalled data; Fig.2A). N50s are represented in Table 1; median N50 across species was 19496bp for R9.4.1 hac, 16002bp for R10.3, 20976bp for R10.4 (all) and 16425bp for R10.4 duplex reads.

**Figure 2.**
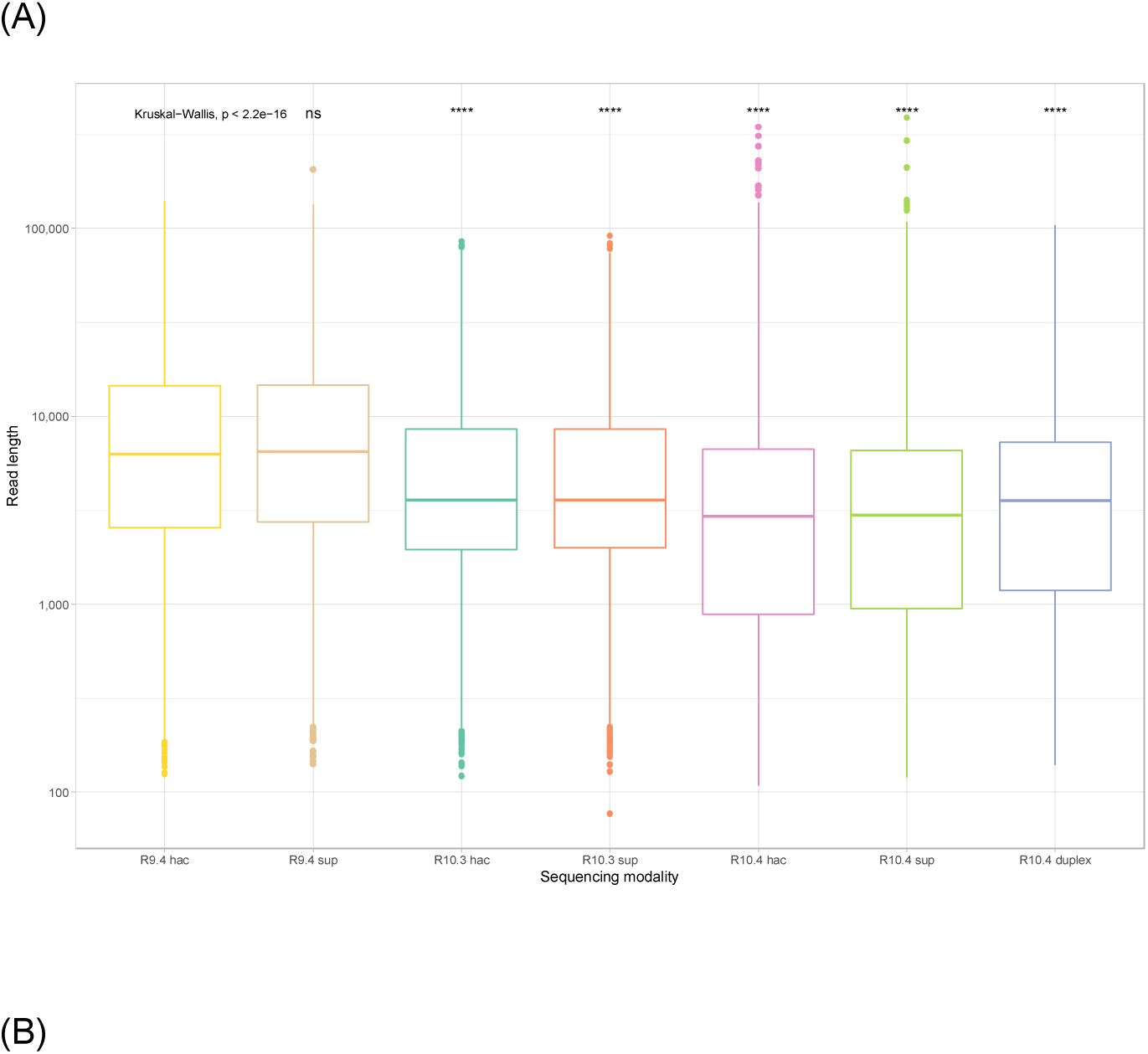
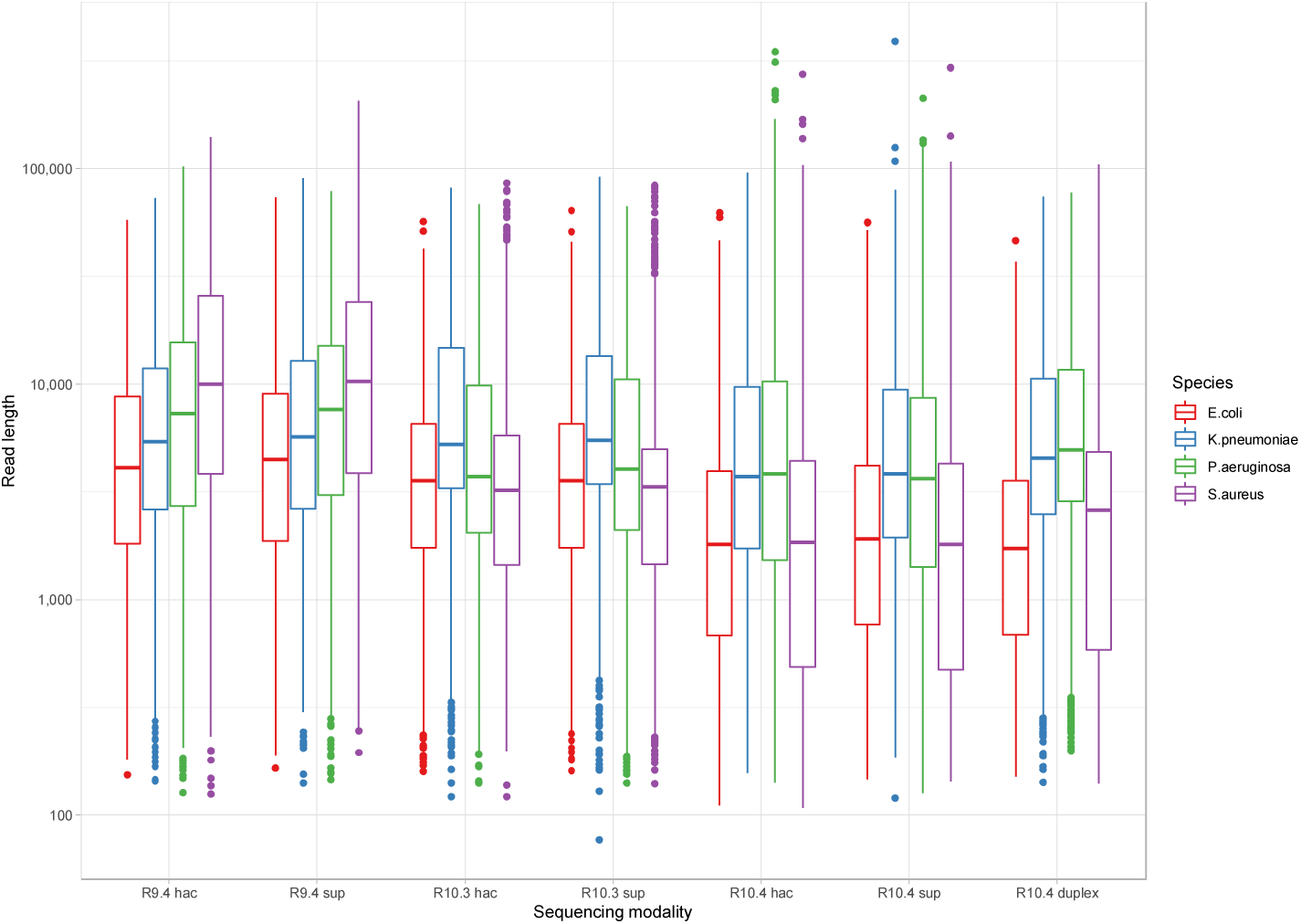
Read length distributions by (A) modality and (B) by modality and species. Boxplots reflect median (central line) and IQR (box hinges) values, whiskers the smallest and largest values 1.5*IQR, and dots the outlying points beyond these ranges. Note the y-axis is a log-scale. Median differences in read length were significant across the whole dataset (Kruskal-wallis, p<0.001); other significance values represent comparisons with the average read length for R9.4 hac as the reference category (“ns” - not significant, “****” - p<0.001).

### Duplex reads

The median proportion of duplex reads across the four unplexed, single-extract R10.4 runs was 4.5% (3.8% for *E. coli*, 6.1% for *K. pneumoniae*, 4.5% for *P. aeruginosa*, and 4.5% for *S. aureus*). For the multiplexed R10.4 run for each species these proportions were 2.3%, 5.4%, 6.0% and 4.7%.

### Raw read accuracy by sequencing modality and species

Raw read accuracy (% identity when mapped to the reference) for a subsample of 1000 reads by sequencing data type/process (i.e. “sequencing modality”) and species was highest (as expected) for Illumina reads (modal accuracy: 100.0%), followed by R10.4 duplex reads basecalled with the sup model (modal accuracy: 99.9%); modal accuracies for all the other approaches were >97.0% (Fig.3). Sup basecalling improved modal accuracy for R10.4 reads, but not R10.3 or R9.4 reads; multiplexing had no impact (Fig.3). Median and modal accuracies for each sequencing modality by species are detailed in Table S6.

**Figure 3.**
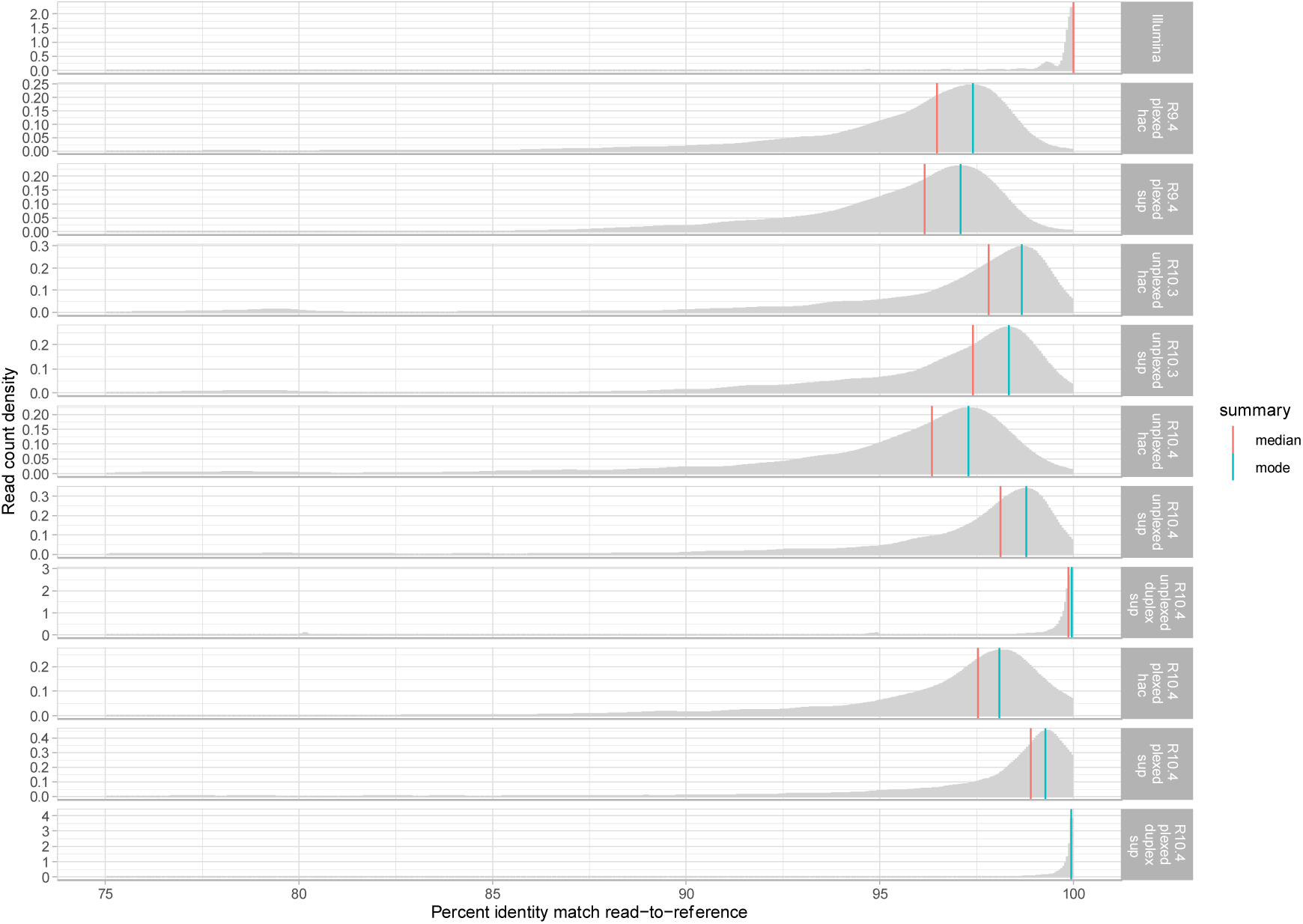
Median and modal raw read accuracy (% identity when reads are mapped to the Illumina-corrected reference) for each of the major nanopore sequencing sequencing modalities, flowcells/kit and basecalling combinations. Reads matching to the reference with <75% identity have been excluded. Complete details summarising all accuracies across all modality, flowcell/kit and basecalling combinations, and stratified by species are represented in Supplementary Table S6.

In terms of insertions and deletions with respect to the reference, for long-read modalities R10.4 sup called duplex data performed best (Fig.4A, 4B). The median number of insertions observed per read was 0.94, 0.45, 0.37 and 0.0 for R9.4 hac, R10.3 hac and R10.4 sup and R10.4 sup duplex respectively (two-sample Wilcoxon test for each versus R9.4 hac as the reference category; all p<0.001), and for deletions 1.31, 0.73, 0.63 and 0.10 respectively (two-sample Wilcoxon test for each versus R9.4 hac as the reference category; all p<0.001).

**Figure 4.**
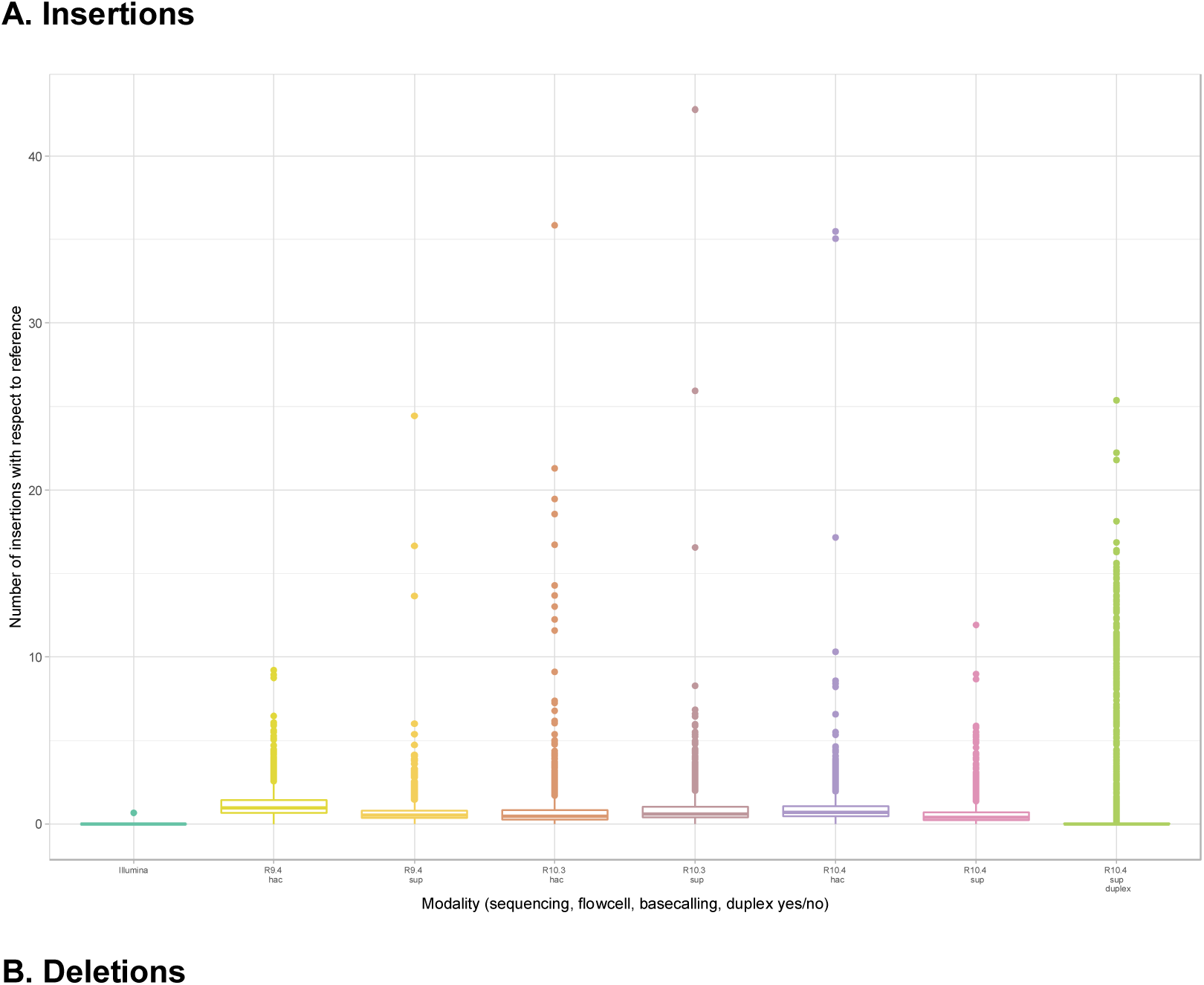
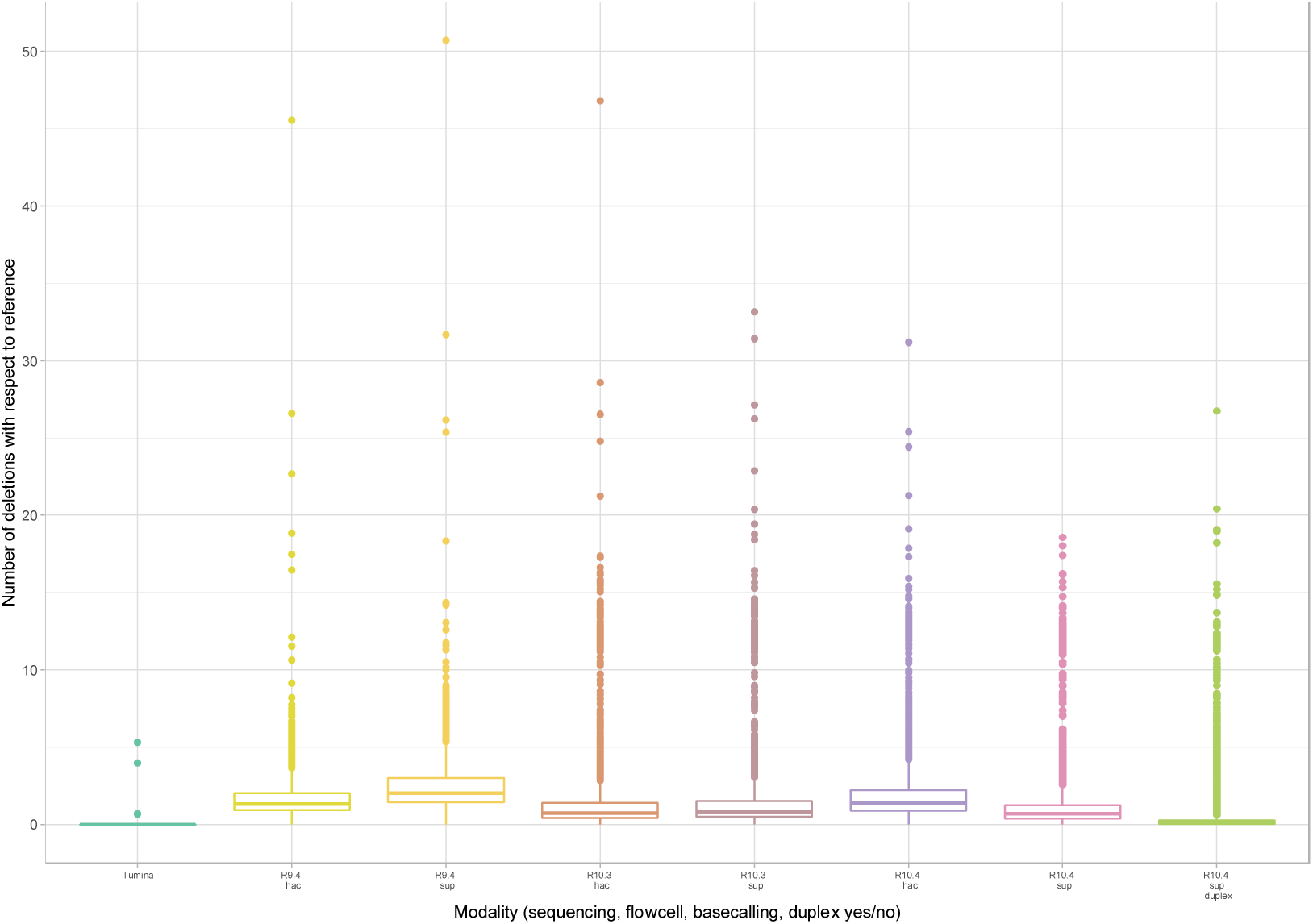
Number of insertions (panel A) and deletions (panel B) amongst reads mapped to the Illumina-corrected reference for all sequencing modalities.

### Assembly accuracy with respect to number of expected contigs in the reference sequences and reference sequence size

We evaluated the capacity of each sequencing approach to accurately reconstruct (i) the number of known contigs present in each reference isolate, and (ii) what percentage of the Illumina-corrected reference was covered. All isolates contained single chromosomes only, except the *K. pneumoniae* reference, which contained a chromosome and five plasmids ranging in size from 3478-175879bp (Table 2).

**Table 2.** Number of unique contigs by sequencing. **modality and bacterial species.** Using the complete data available (i.e. no subsampling). The first number in each cell represents the number of contigs assembled and matching to the Illumina-corrected reference using dnadiff, the total number of contigs assembled is shown in curved brackets, and the proportion of the reference chromosomal contig covered in square brackets. For *E. coli*, *P. aeruginosa*, *S. aureus* the total number of expected contigs is 1, for *K. pneumoniae* 1 chromosome + 5 plasmids. Orange shading shows absent contigs, and/or incomplete assembly (n >1 contig matching to reference), and/or extra contigs not matching to reference. Green shaded cells denote complete singular contigs which reflect the reference DNA content at 100+/-1%. “-“ denotes no relevant contig assembled.

| Sequencing modality | Assembler | Plexed Y/N | *E. coli*, chromosome (1)<br>5231428bp<br>51% GC | *P. aeruginosa*, chromosome (1)<br>6264404bp<br>66.2%GC | *S. aureus*, chromosome (1)<br>2902619bp<br>32.8%GC | *K. pneumoniae*, chromosome (1)<br>5315120bp<br>57.0%GC | *K. pneumoniae*, pKPN3 plasmid (1)<br>175879<br>51.7%GC | *K. pneumoniae*, pKPN4 plasmid (1)<br>107576bp<br>53.4%GC | *K. pneumoniae*, pKPN5 plasmid (1)<br>88582bp<br>53.8%GC | *K. pneumoniae*, pKPN6 plasmid (1)<br>4259bp<br>41.4%GC | *K. pneumoniae*, pKPN7 plasmid (1)<br>3478bp<br>45.7%GC |
| --- | --- | --- | --- | --- | --- | --- | --- | --- | --- | --- | --- |
| Illumina | SPAdes | Y | 226 (326) [98.64%] | 115 (152) [99.72%] | 86 (150) [98.42%] | 117 (312) [98.9%] | 41 [79.7%] | 45 [100.72%] | 22 [82.49%] | 1 [102.82%] | 1 [103.45%] |
| R9.4.1 hac+Illumina | Unicycler | Y | 1 (1) [100.09%] | 1 (1) [100.49%] | 1 (1) [100.69%] | 1 (7) [100.1%] | 1 [100%] | 1 [100%] | 1 [100%] | 2 [100%] | 1 [96.15%] |
| R9.4.1 hac | Canu | Y | 1 (1) [100.59%] | 1 (1) [101.29%] | 1 (1) [103.18%] | 1 (24) [100.72%] | 1 [131.55%] | 1 [172.51%] | 1 [100%] | 2 [133.5%] | 1 [100%] |
| R9.4.1 sup | Canu | Y | 1 (1)<br>[100.65%] | 1 (1)<br>[101.18%] | 1 (1)<br>[102.43%] | 1 (7)<br>[100.96%] | 1<br>[129.41%] | 1 [100%] | 1<br>[125.07%] | 1 [100%] | 1 [100%] |
| R10.3 hac | Canu | N | 1 (6)<br>[100.49%] | 1 (2)<br>[101.19%] | 1 (3)<br>[102.78%] | 1 (8)<br>[100.72%] | 1 [100%] | 1 [100%] | 1 [100%] | 1 [100%] | 1 [100%] |
| R10.3 sup | Canu | N | 1 (2)<br>[100.6%] | 1 (3)<br>[101.13%] | 1 (3)<br>[102.85] | 1 (8)<br>[100.74%] | 1 [100%] | 1<br>[138.13%] | 1<br>[143.94%] | 1<br>[192.58%] | 1 [100%] |
| R10.4 hac | Canu | N | 1 (6)<br>[100.62%] | 1 (1)<br>[101.06%] | 1 (2)<br>[103.6%] | 1 (8)<br>[100.8%] | 1<br>[127.89%] | 1<br>[141.81%] | 1 [147.6%] | 1 [100%] | 1<br>[112.85%] |
| R10.4 sup | Canu | N | 1 (3)<br>[100.61%] | 1 (2)<br>[100.1%] | 1 (2)<br>[102.33%] | 1 (7)<br>[100%] | 1<br>[100.94%] | 1 [100%] | 1<br>[145.43%] | 1 [100%] | 1 [100%] |
| R10.4 sup<br>duplex | Canu | N | 4 (42)<br>[100.12%] | 1 (14)<br>[101.5%] | 1 (29)<br>[101.72%] | 1 (41)<br>[100.1%] | 1 [100%] | 3<br>[130.98%] | 1<br>[149.19%] | 1 [100%] | 1 [100%] |
| R10.4 hac | Canu | Y | 1 (2)<br>[100.42%] | 1 (1)<br>[100.87%] | 1 (1)<br>[102.01%] | 1 (7)<br>[100.82%] | 1<br>[124.01%] | 1 [100%] | 1<br>[142.89%] | 1 [100%] | 1 [100%] |
| R10.4 sup | Canu | Y | 1 (1)<br>[100.48%] | 1 (1)<br>[100.17%] | 1 (1)<br>[102.29%] | 1 (12)<br>[100.61%] | 1 [117.3%] | 2<br>[134.06%] | 1<br>[137.68%] | 1 [100%] | 1 [100%] |
| R10.4 sup<br>duplex | Canu | Y | -* | 23 (25)<br>[99.2%] | -* | 15 (25)<br>[99.2%] | 1 [80.39%] | 2 [97.73%] | 1<br>[112.86%] | 1 [94.01%] | 1 [100%] |
| R9.4.1 hac | Flye | Y | 1 (1)<br>[100.09%] | 1 (1)<br>[100.14%] | 1 (1)<br>[100.7%] | 1 (4)<br>[100.11%] | 1 [75.7%] | 1<br>[103.68%] | 1 [100%] | - | - |
| R9.4.1 sup | Flye | Y | 1 (1)<br>[100.09%] | 1 (1)<br>[100.47%] | 1 (1)<br>[100%] | 1 (5)<br>[100.1%] | 1 [100%] | 1 [99.99%] | 1 [100%] | - | - |
| R10.3 hac | Flye | N | 1 (1)<br>[100.10%] | 1 (1)<br>[100.44%] | 1 (1)<br>[100.69%] | 1 (4)<br>[100.1%] | 1 [95.83%] | 1 [73.13%] | 1 [100%] | - | - |
| R10.3 sup | Flye | N | 1 (1)<br>[100.09%] | 1 (1)<br>[100.44%] | 1 (1)<br>[100.69%] | 1 (4)<br>[100.1%] | 1 [100%] | 1 [100%] | 1 [100%] | - | - |
| R10.4 hac | Flye | N | 1 (1)<br>[100.10%] | 1 (1)<br>[100.11%] | 1 (1)<br>[100.69%] | 1 (4)<br>[100.1%] | 1 [100%] | 1 [100%] | 1 [100%] | - | - |
| R10.4 sup | Flye | N | 1 (1)<br>[100.09%] | 1 (1)<br>[100.4%] | 1 (1)<br>[100.69%] | 1 (5)<br>[100.1%] | 1 [100%] | 1 [98.92%] | 1 [99.99%] | - | - |
| R10.4 sup<br>duplex | Flye | N | 1 (3)<br>[100.10%] | 1 (1)<br>[100.8%] | 1 (2)<br>[100.69%] | 1 (7)<br>[100.21%] | 1 [94.27%] | 1<br>[102.93%] | 1<br>[101.54%] | - | 1 [100%] |
| R10.4 hac | Flye | Y | 1 (1)<br>[100.10%] | 1 (1)<br>[100.16%] | 1 (1)<br>[100.69%] | 1 (5)<br>[100.1%] | 1 [100%] | 1 [100%] | 1 [100%] | - | - |
| R10.4 sup | Flye | Y | 1 (1)<br>[100.00%] | 1 (1)<br>[100.48%] | 1 (1)<br>[100.69%] | 1 (5)<br>[100.11%] | 1 [100%] | 1 [100%] | 1 [100%] | - | - |
| R10.4 sup<br>duplex | Flye | Y | 37 (38)<br>[8.47%] | 1 (5)<br>[100.48%] | 25 (25)<br>[83.64%] | 1 (4)<br>[100.4%] | 1 [84.71%] | 1<br>[105.27%] | 1 [100%] | - | - |

Approaches using all the data and Unicycler or Flye largely generated single chromosomal contigs, except those using R10.4 duplex reads only, particularly for multiplexed extracts, likely because these reads were insufficient to cover the whole genome (Table 2; Fig.S5A). Illumina-only assemblies generated much larger numbers of contigs as expected (Table 2). Using all the data, single *K. pneumoniae* plasmid contigs were mostly obtained using any of the long-read data and Flye, or hybrid assembly with Unicycler (Table 2, Fig.S5B). Using all the data, Flye long-read only assemblies largely all missed the two smallest plasmids (Table 2, Fig.S5B).

Sub-sampling the data to 10x, 20x, 30x, 40x, 50x or 100x depth had variable effect - for the most part single chromosomal contigs were assembled using long-reads only with >20x depth; Unicycler could mostly be used with 10x long-read depth (Fig.S5A). The same effect was seen for plasmids, except Flye struggled to reliably assemble the two largest plasmids into single contigs with lower sequencing depths (Fig.S5B). Canu assemblies failed with 10x sub-sampling, as expected given the default cut-offs.

For chromosomes, Canu long-read only assemblies tended to over-assemble structures (i.e. reference coverage >100%, Fig.5A) whilst Illumina-only assemblies under-assembled structures. Reference coverage % for Unicycler hybrid (R9.4+Illumina) was largely unaffected by sub-sampling the data to 10x, 20x, 30x, 40x, 50x or 100x (Fig.5A). For plasmids, Canu assembly again largely over-assembled the structures; Unicycler hybrid (R9.4+Illumina) assembly was the only approach which consistently assembled all plasmids at near 100% reference coverage across all sub-sampling depths (Fig.5B).

**Figure 5.**
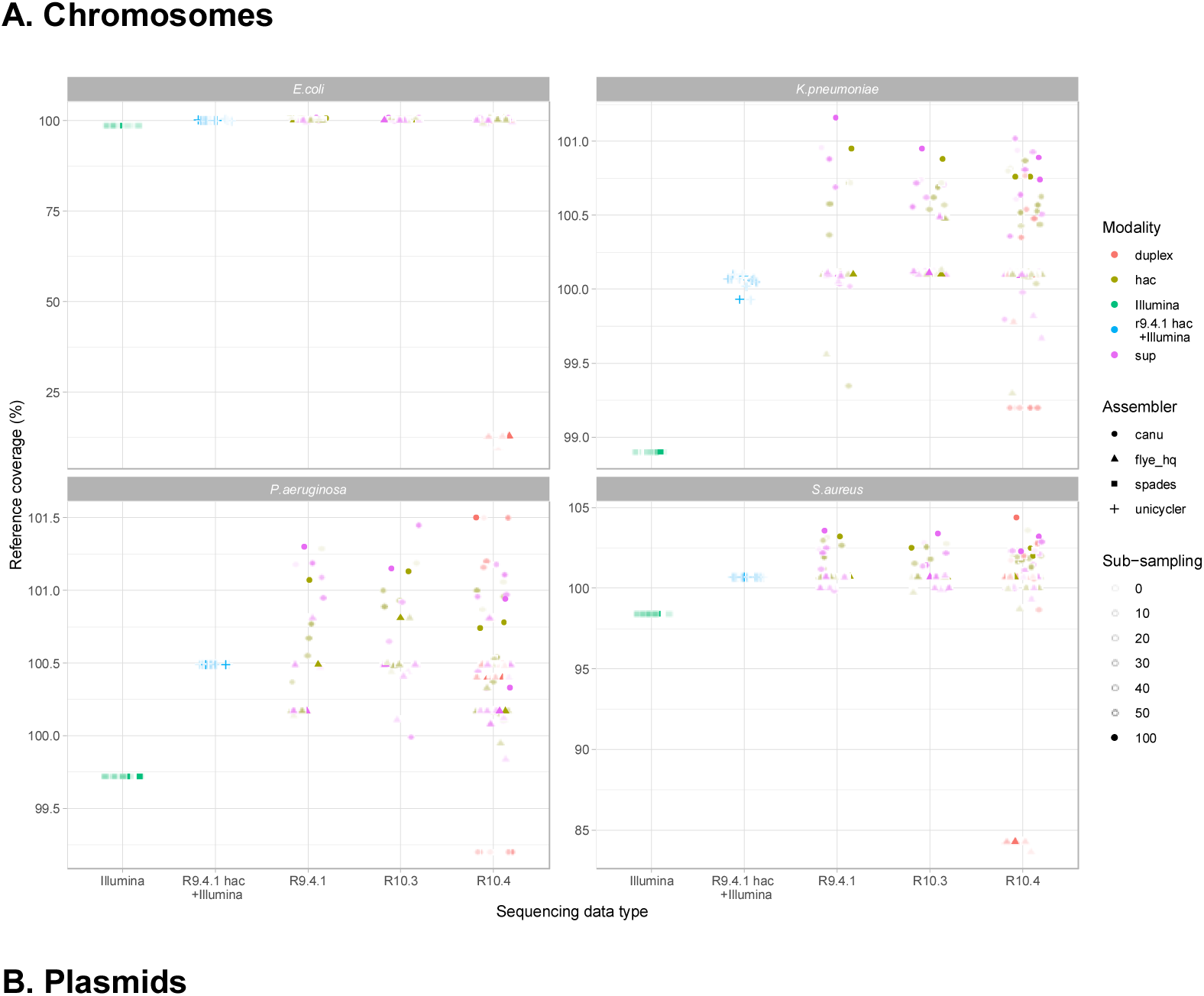
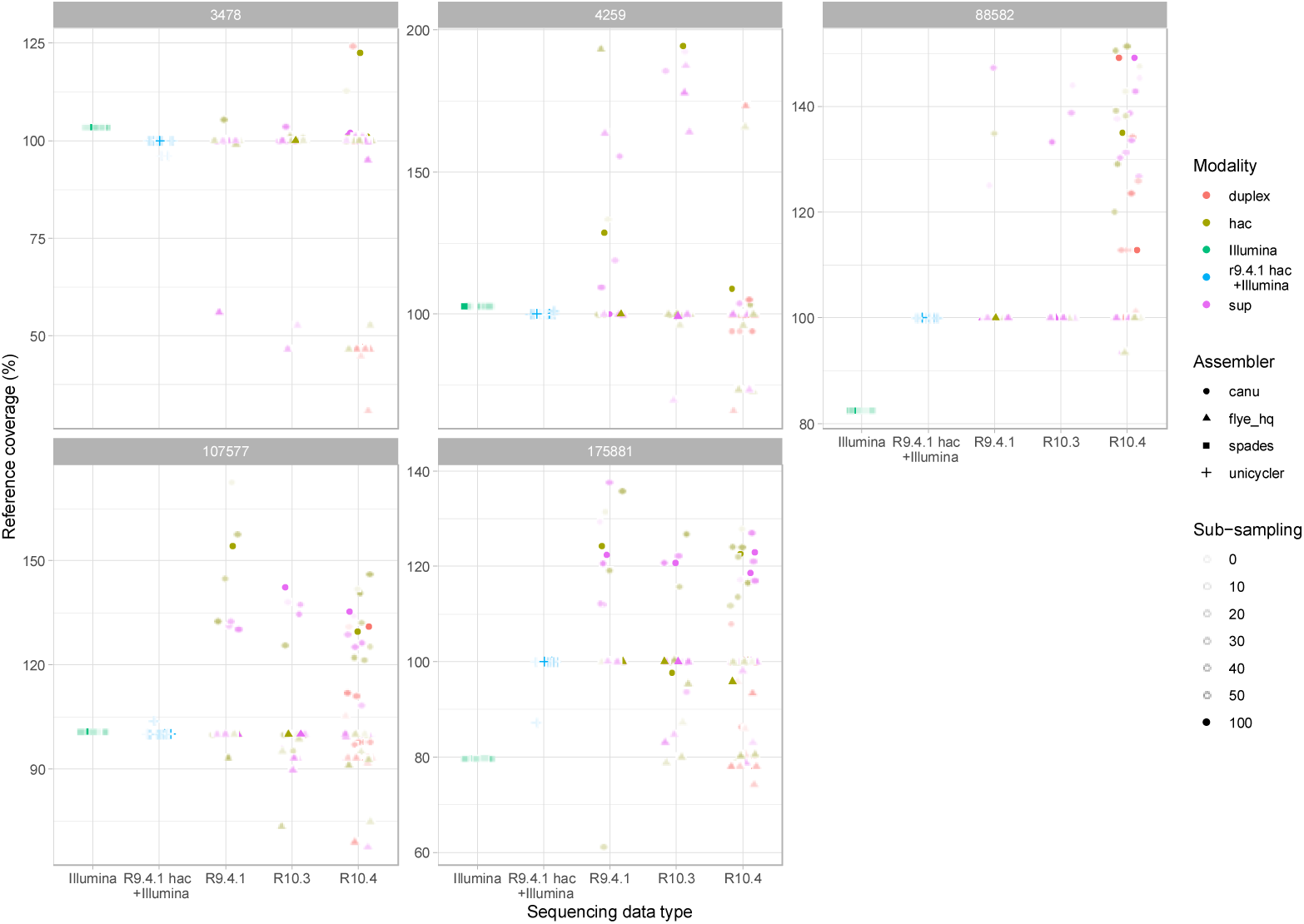
Assembly reference coverage percentage (%) by sequencing modality, assembler and species. Panel A represents the data for chromosomes and panel B evaluations for the five plasmids known to occur in the *K. pneumoniae* reference strain (labeled by their lengths in bp).

### Assembly accuracy with respect to insertions, deletions and nucleotide-level mismatches

For each sequencing and assembly modality the number of indels and nucleotide-level mismatches (SNPs) were evaluated by species (Figs.6A, 6B) and overall (Table S7). The impact of sub-sampling and relevance of long-read sequencing depth was also considered (Fig.7).

Overall, SPAdes assemblies had the fewest indels (0.02 indels/100kb), followed by Medaka-polished Flye-assembled R10.4 sup basecalled/duplex reads (0.18 indels/100kb), Medaka-polished Flye-assembled R10.4 sup basecalled data (0.41 indels/100kb), Medaka-polished Flye-assembled R10.3 hac basecalled data (for 3 rounds of polishing: 0.44 indels/100kb) and Unicycler assemblies (0.56 indels/100kb) (Table S7). There were apparent species-specific differences, with the *E. coli* reference proving the most challenging to assemble accurately (Fig.6A). The improvements in the indel error rates of R9.4 or R10.4 Flye assemblies polished with 2 or 3 rounds of Medaka versus 1 round were negligible; however, additional rounds of polishing improved indel errors in R10.3 hac basecalled assemblies (Fig.6A, Fig.S6, Table S7).

**Figure 6.**
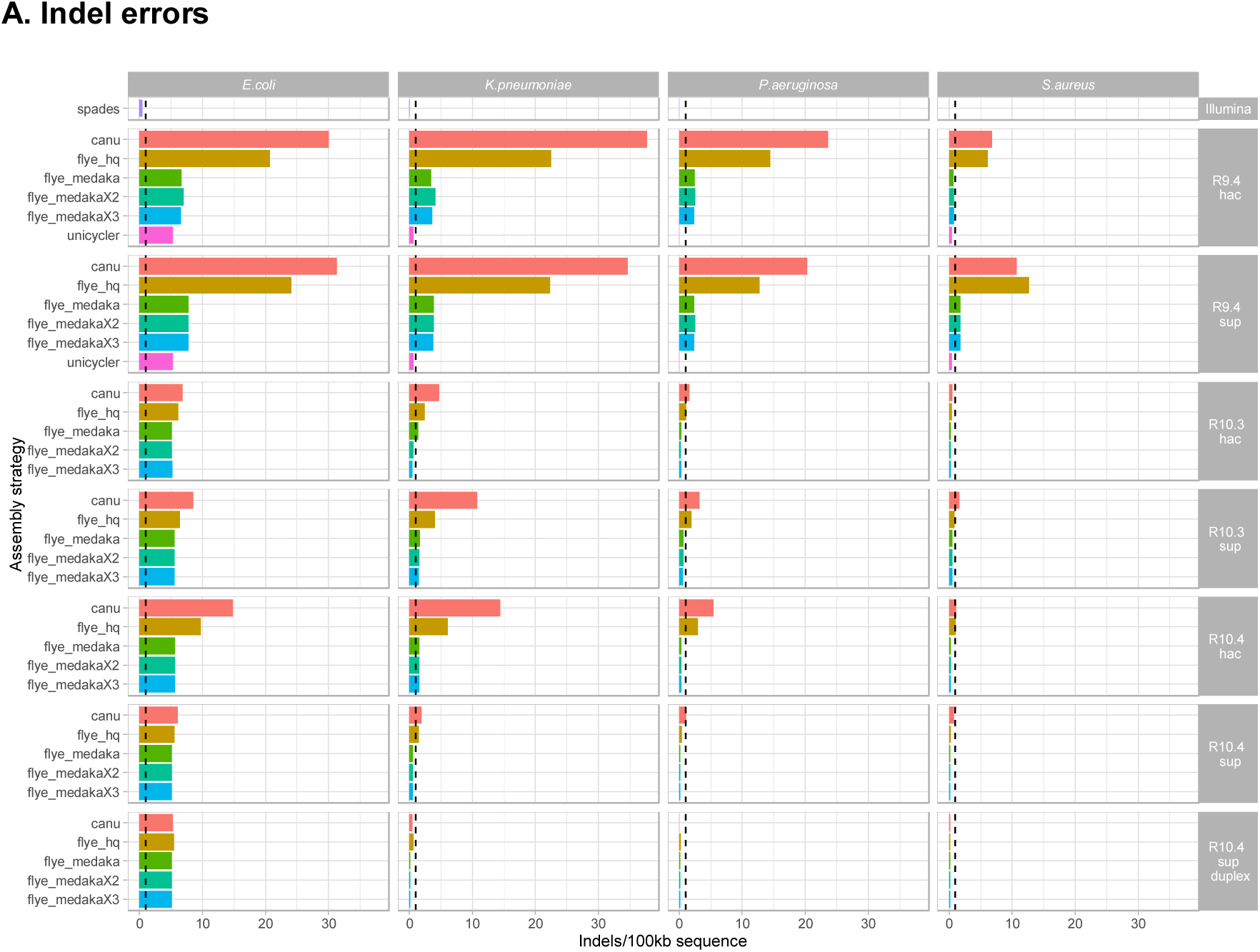
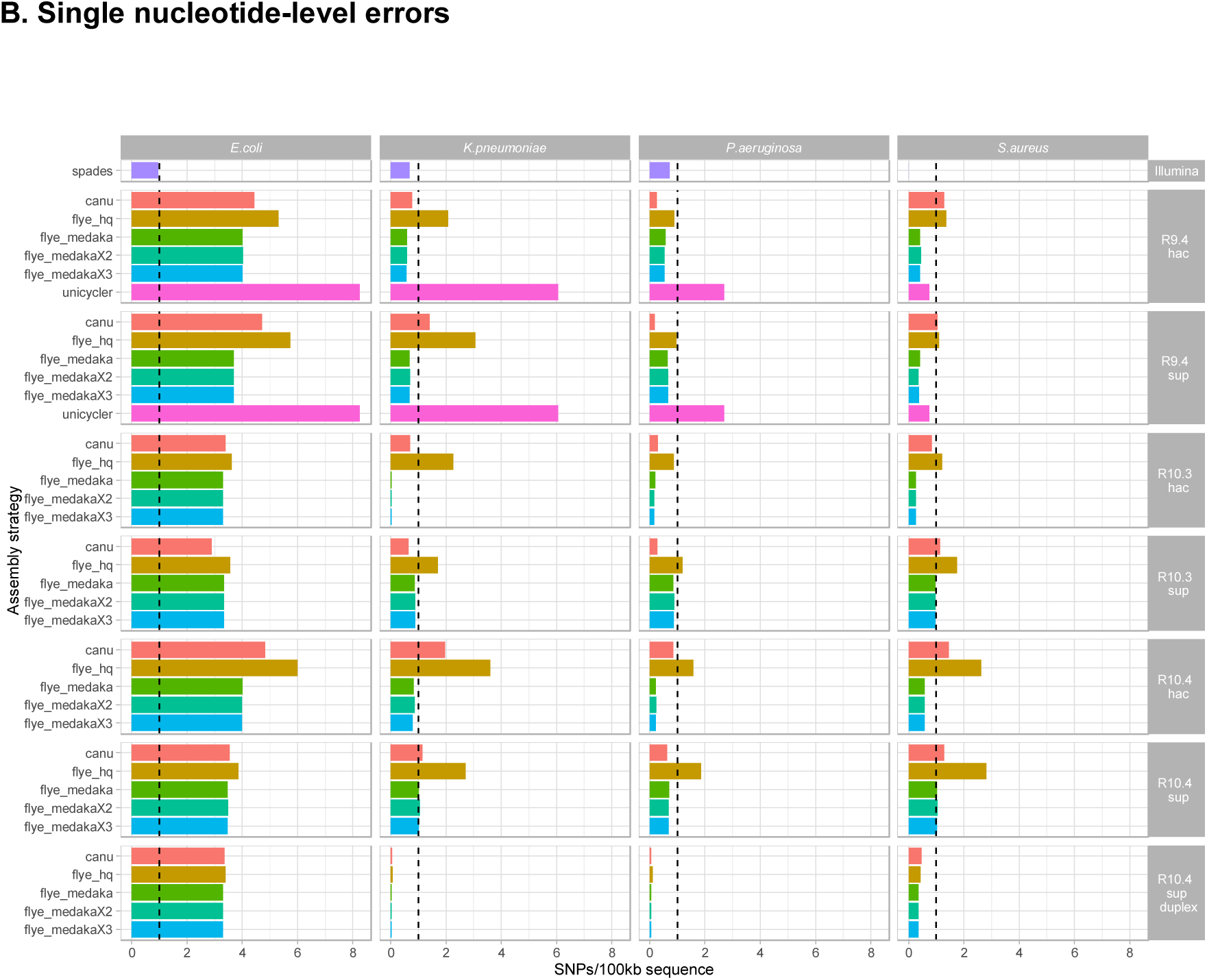
Assembly accuracy by sequencing modality, assembly strategy and species. Accuracy evaluated on the basis of contig comparisons to Illumina-corrected references using dnadiff, for (A) Indels, and (B) SNPs. NB - SPAdes was only used on Illumina data, and Unicycler hybrid assembly was only performed on R9.4.1+Illumina data. For R10.4, data presented are those from unplexed runs. Dashed black vertical line indicates a threshold of 1 error/100kb.

Similar trends were observed overall for SNPs, with the lowest error rates (0.21 SNPs/100 kb of sequence) observed for multiply-Medaka-polished Flye-assembled R10.3 hac basecalled data, or singly-Medaka-polished Flye assembled R10.4 sup basecalled/duplexed data (0.21 SNPs/100kb of sequence) (Fig.6B, Table S7). SNP error rates for Unicycler assemblies however were higher than for the other optimised assembly modalities (4.38 SNPs/100kb) (Tables S7). Polishing Flye assemblies with Medaka improved SNP error rates over unpolished assemblies, but there were no obvious benefits of multiple rounds of polishing (Fig.6B, Fig.S6). Again, species-specific differences were observed, with the *E. coli* reference the most challenging to assemble (Fig.6B).

Error rates for Unicycler assemblies were largely consistent at all long-read sequencing depths from 10X to up to strategies using all the data; error rates for long-read-only assemblies were optimised when coverage was ≥20X (Fig.7).

**Figure 7.**
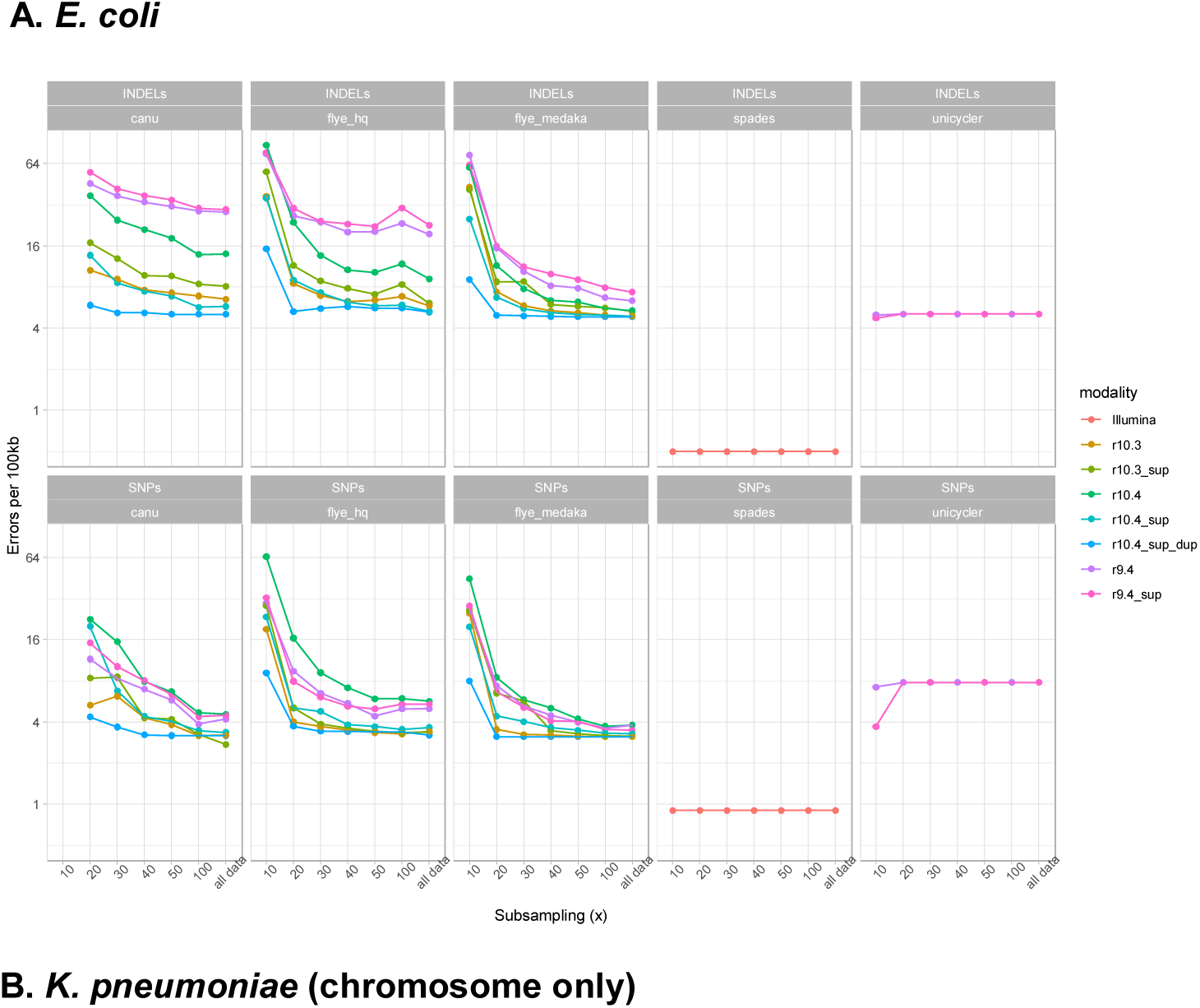
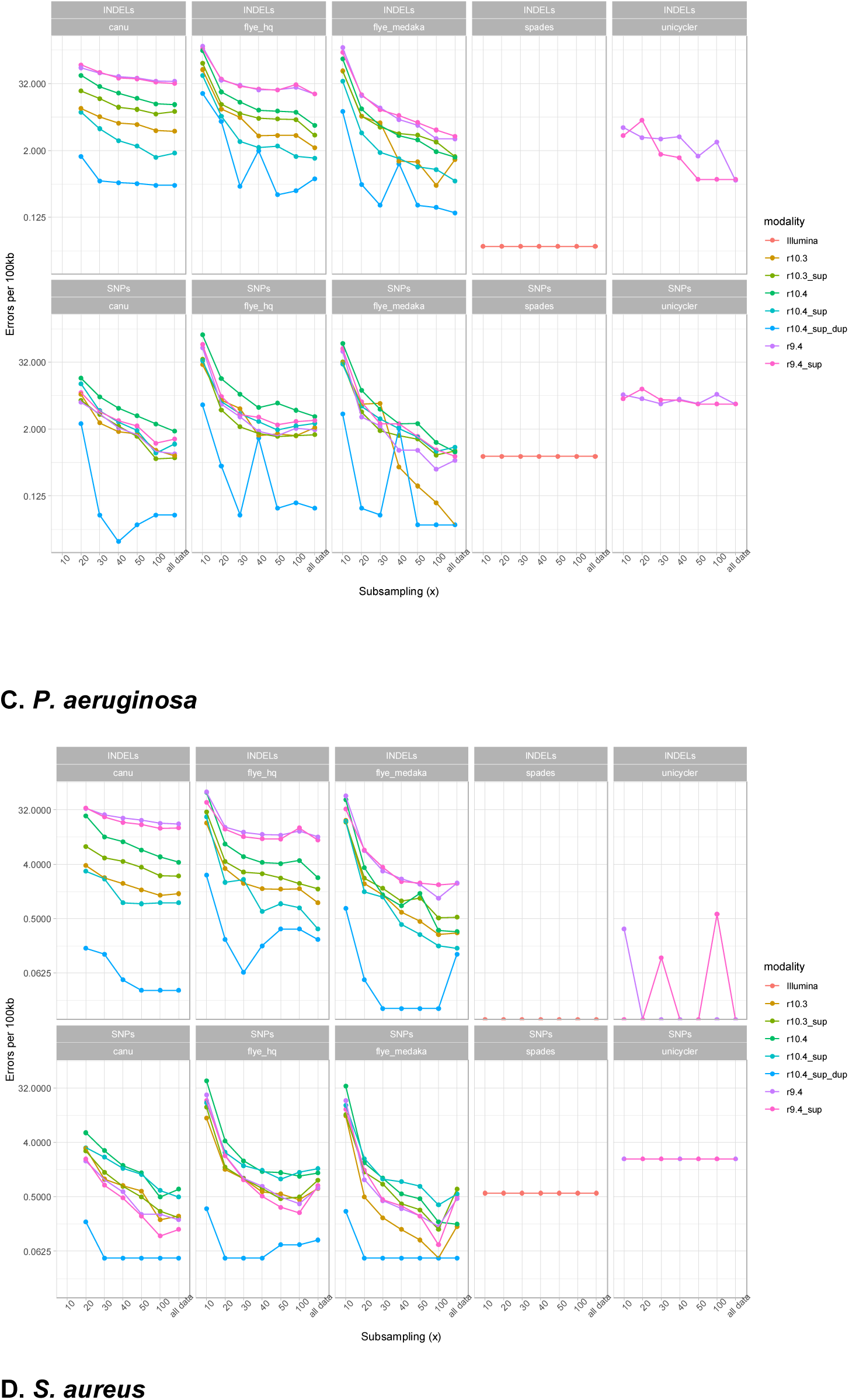
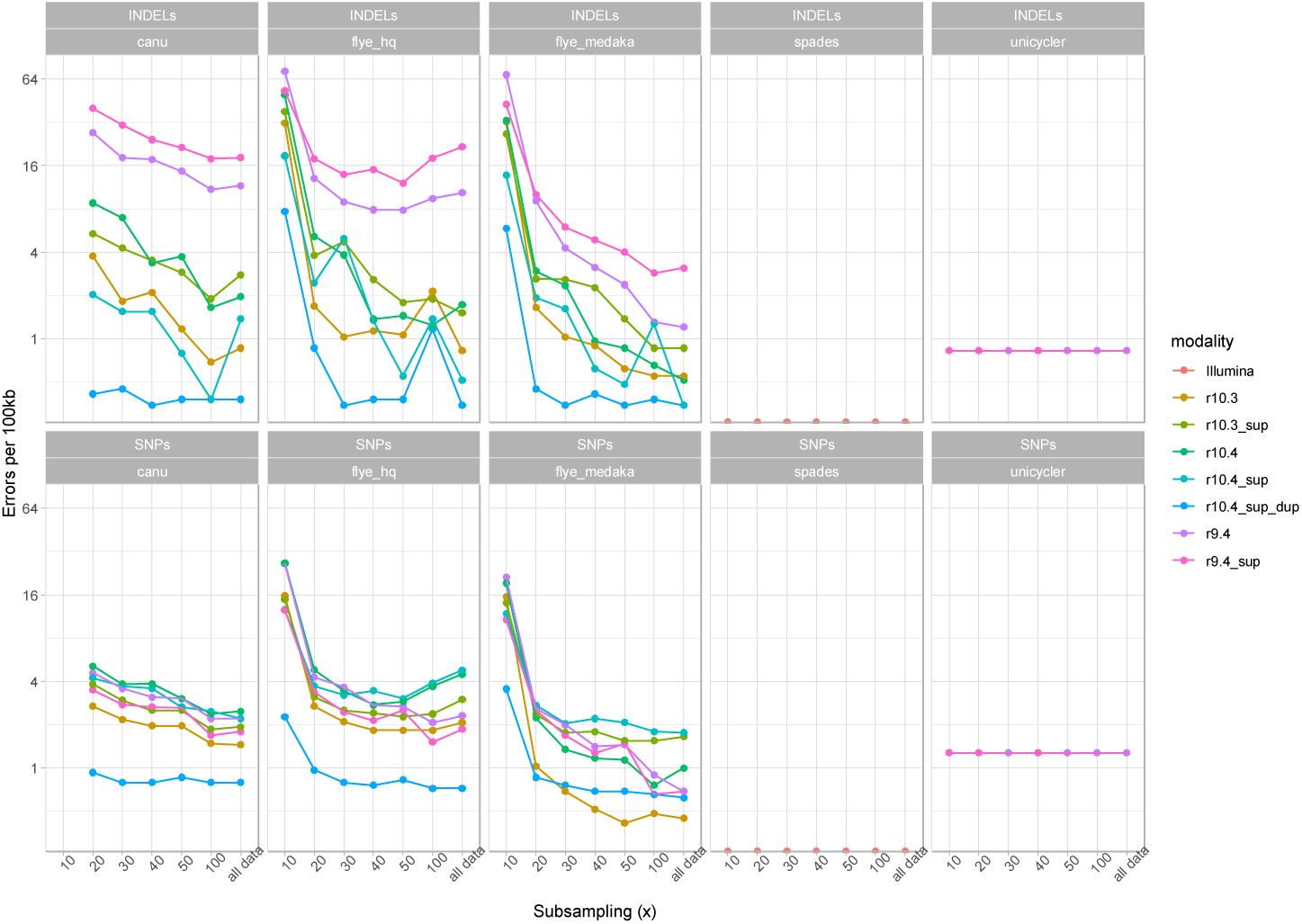
Impact of subsampling of long-read datasets on assembly accuracy. Presented here by species for Indels (top panels), and SNPs (lower panels). For ease of representation, only data for Flye assemblies polished with 1 round of Medaka are shown, as the effects of additional polishing was shown to be marginal for most modalities (Fig.S6, Table S7). Data for 10x long-read coverage is not omitted for Canu assemblies as this coverage was considered too low for default settings and was unlikely to improve results.

### Assembly accuracy with respect to coding sequence content

Coding sequence content was most accurately recovered using Flye-assembled sup basecalled R10.4 duplex data and hybrid assembly (Fig.8; missing between 9-32 (∼0.25-0.75%) of coding sequences across species). Long-read only assembly with R9.4 data missed up to 10-15% of coding sequences (data not plotted in Fig.8). Notably, the duplex datasets from the unplexed 10.4 runs were used, as from multiplexed runs the duplex yields were insufficient to facilitate assembly in most cases (Table 2).

**Figure 8.**
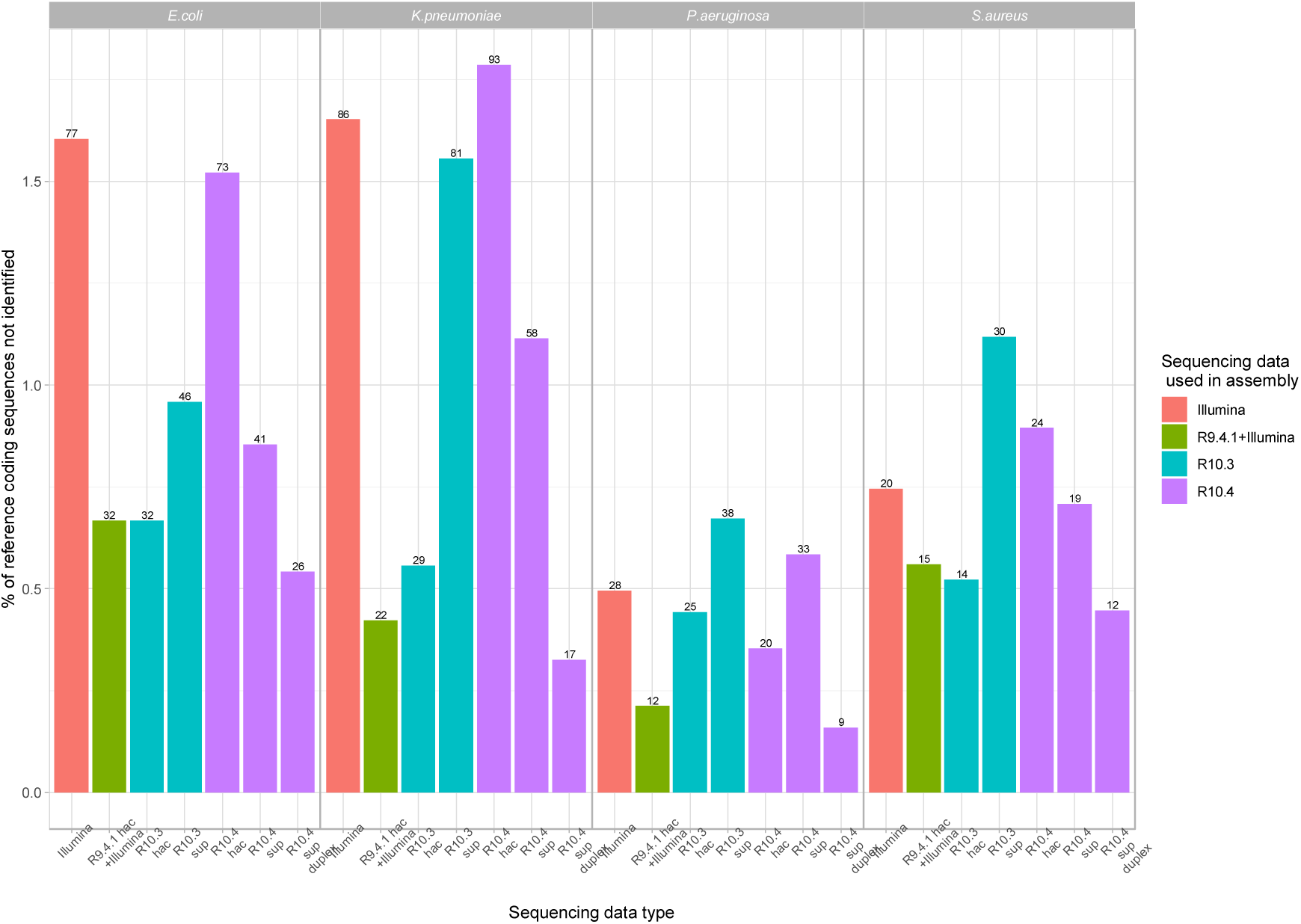
Coding sequence (CDS) recovery on the basis of exact CDS (amino acid sequence) matches with respect to the Prokka-annotated Illumina-corrected reference (chromosome+all plasmids for *K. pneumoniae*). Plot shows the percentage of reference coding sequences missed by each modality. For long-read data only Flye assemblies with one round of polishing with Medaka are shown shown; for R10.3 and R10.4 datasets these were from non-multiplexed evaluations (i.e. only single extracts per flowcell). For Unicycler, the assembly using R.9.4 hac+Illumina data is shown. The total number of coding sequences missed by each approach is shown as a number at the top of each bar.

## 8. Discussion

In this pragmatic study evaluating the impact of different nanopore sequencing flowcells and chemistries on the capacity to fully reconstruct genomes of four commonly studied bacteria, we have shown that sup basecalled R10.4/Kit12 data and sup called duplex data have read- and assembly-level accuracies that would enable these to be effectively used for the reconstruction of bacterial genomes without requiring Illumina data to generate hybrids. However, hybrid assembly (Illumina+9.4.1 hac data) remains the most robust approach in terms of contig (both chromosomes and plasmids) and CDS recovery without over-assembly, and facilitates the multiplexing of large numbers of isolates per flowcell, given that in this and at least one other study(3), ≤10x long-read depth is required for the accurate reconstruction of chromosomes and plasmids by combining R9.4.1 and Illumina data using Unicycler. Highly accurate long-read only assembly and genome reconstructions was optimized by generating duplex reads, which in our hands made up a small proportion of the output (<10%); as such, it would come at a significant cost per isolate as a result of being able to only generate data for 1-2 isolates per flowcell. Very approximate costs per genome therefore for hybrid assembly versus duplex/long-read-only assembly would be £50-70/genome versus £300-600/genome.

Although barcoding up to 96 isolates has recently been enabled for the R10.4/Kit12 combination, the data yields per flowcell (∼4Gb) would likely preclude viable assembly for 96 *E. coli* isolates with a typical genome size of ∼5Mb (would give <8x coverage). There is also a current requirement to use a ligation-based library preparation, which lengthens the processing time, and may impact on plasmid recovery(6). We observed issues with recovering small plasmids (<5kb) using Flye in this study although both of these small plasmids could be reliably recovered in Canu assemblies; consistent with this a previous evaluation has shown that 8-15% of small plasmids are not recovered using these long-read-only assemblers(12). Similarly, as shown in this study and in other work(12), the basic Canu workflow ‘over-assembles’ the data, and contigs require trimming of overlaps in order to recreate accurate, single, circularized structures. We observed some apparent species-specific differences, suggesting that assemblers are more challenged in accurately reconstructing certain genomes; these differences, as well as differences related to genome length and the impact on long-read sequencing depth may be important to consider in study design.

There are currently few other published studies on the performance of R10.4/Kit12 for bacterial analyses. We found only one preprint investigating its use on a mock microbial community (7 bacterial species and 1 fungal species) which found similar modal accuracy scores of 99% using sup basecalling, and a requirement of 40x to be able to reliably assemble a genome(20). Their hypothesis was that improved read accuracies were due to an improved ability to call homopolymers, which we did not investigate in this manuscript. It was unclear what proportion of reads they characterized as duplex reads.

There are several limitations of our study. We have not exhaustively investigated all possible approaches to genome assembly, but rather taken a pragmatic approach in assembling the data with several commonly used assemblers, without additional bespoke management or combination of workflows; the data are however available for other researchers to trial different approaches. We had low duplex read yields compared with those reported by ONT (up to 30-40% per flowcell); further optimization is needed to see if these can be achieved. We have investigated only a limited number of isolates and plasmids, but these represent a range of %GC and sizes, and are likely to reflect genetic content more widely in other species; we have not generated replicate datasets. Similarly, because we only investigated one isolate per species, it may be that the differences observed are not generalisable or are strain and not species-specific; this would be interesting future work. Improvements and upgrades to nanopore flowcells, chemistries and basecallers occur regularly and nanopore will be releasing the R10.4.1 flowcell and kit14 chemistries later in 2022 which may further optimise the quality of long-read only outputs.

In summary, the combination of R10.4/Kit12 flowcells/chemistries look very promising for highly accurate, long-read only bacterial genome assembly; however, this requires superior accuracy basecalling, and is optimised by the generation of duplex reads, which currently make up only a small proportion of sequencing yield. In addition, for large-scale projects to fully reconstruct 100s-1000s of bacterial isolates, hybrid assembly, multiplexing and the use of flowcells/chemistries that support rapid barcoding are currently better suited for higher throughput and are more cost-effective per reconstructed genome. The optimal strategy in any given context will depend on the specific use case and resources available, and may evolve rapidly over short timescales.

## 9. Author statements

### 9.1 Authors and contributors

NK, NSa, DWC and NS designed the study. NK and GR performed the laboratory experiments and sequencing. NSa performed the bioinformatics analysis. NSt generated the data visualisations. NSt, NSa and NK wrote the first draft. All authors reviewed and approved the final draft.

### 9.4 Ethical approval

Not applicable.

### 9.5 Consent for publication

Not applicable.

## Acknowledgements

We are grateful to Dr Celiq Souque and Prof Craig Maclean at the Department of Zoology, University of Oxford, for supplying the *Pseudomonas aeruginosa* PAO1 strain. We are also grateful for feedback from the Twitter community following the release of this manuscript as a preprint.

## References

1. Van Goethem N, Descamps T, Devleesschauwer B, Roosens NHC, Boon NAM, Van Oyen H, et al. Status and potential of bacterial genomics for public health practice: a scoping review. Implementation science : IS. 2019;14(1):79.

2. Shaw LP, Chau KK, Kavanagh J, AbuOun M, Stubberfield E, Gweon HS, et al. Niche and local geography shape the pangenome of wastewater- and livestock-associated Enterobacteriaceae. Science advances. 2021;7(15).

3. Arredondo-Alonso S, Pöntinen AK, Cléon F, Gladstone RA, Schürch AC, Johnsen PJ, et al. A high-throughput multiplexing and selection strategy to complete bacterial genomes. GigaScience. 2021;10(12).

4. Lipworth S, Pickford H, Sanderson N, Chau KK, Kavanagh J, Barker L, et al. Optimized use of Oxford Nanopore flowcells for hybrid assemblies. Microb Genom. 2020;6(11).

5. Oxford Nanopore Technologies. https://nanoporetech.com/about-us/news/r103-newest-nanopore-high-accuracy-nanopore-sequencing-now-available-store; last accessed: 07/Apr/2022.

6. Wick RR, Judd LM, Wyres KL, Holt KE. Recovery of small plasmid sequences via Oxford Nanopore sequencing. Microb Genom. 2021;7(8).

7. Benton M. 2021. Nanopore Guppy GPU basecalling on Windows using WSL2https://hackmd.io/@Miles/rkYKDHPsO. Blog post, last accessed: 07/Apr/2022.

8. Shen W, Le S, Li Y, Hu F. SeqKit: A Cross-Platform and Ultrafast Toolkit for FASTA/Q File Manipulation. PLoS One. 2016;11(10):e0163962.

9. Hall MB. Rasusa: Randomly subsample sequencing reads to a specified coverage. The Journal of Open Source Software.

10. Koren S, Walenz BP, Berlin K, Miller JR, Bergman NH, Phillippy AM. Canu: scalable and accurate long-read assembly via adaptive k-mer weighting and repeat separation. Genome Res. 2017;27(5):722–36.

11. Kolmogorov M, Yuan J, Lin Y, Pevzner PA. Assembly of long, error-prone reads using repeat graphs. Nature biotechnology. 2019;37(5):540–6.

12. Wick RR, Holt KE. Benchmarking of long-read assemblers for prokaryote whole genome sequencing. F1000Research. 2019;8:2138.

13. Wick RR, Judd LM, Gorrie CL, Holt KE. Unicycler: Resolving bacterial genome assemblies from short and long sequencing reads. PLoS Comput Biol. 2017;13(6):e1005595.

14. Prjibelski A, Antipov D, Meleshko D, Lapidus A, Korobeynikov A. Using SPAdes De Novo Assembler. Curr Protoc Bioinformatics. 2020;70(1):e102.

15. De Maio N, Shaw LP, Hubbard A, George S, Sanderson ND, Swann J, et al. Comparison of long-read sequencing technologies in the hybrid assembly of complex bacterial genomes. Microb Genom. 2019.

16. Klockgether J, Munder A, Neugebauer J, Davenport CF, Stanke F, Larbig KD, et al. Genome diversity of Pseudomonas aeruginosa PAO1 laboratory strains. J Bacteriol. 2010;192(4):1113–21.

17. Chandler CE, Horspool AM, Hill PJ, Wozniak DJ, Schertzer JW, Rasko DA, et al. Genomic and Phenotypic Diversity among Ten Laboratory Isolates of Pseudomonas aeruginosa PAO1. J Bacteriol. 2019;201(5).

18. Kurtz S, Phillippy A, Delcher AL, Smoot M, Shumway M, Antonescu C, et al. Versatile and open software for comparing large genomes. Genome Biol. 2004;5(2):R12.

19. Seemann T. Prokka: rapid prokaryotic genome annotation. Bioinformatics. 2014;30(14):2068–9.

20. Sereika MK, R.H.; Karst, S.M.; Michaelsen, T.Y.; Soresnes, E.A.; Wollenberg, R.D.; Albertsen, M. Oxford Nanopore R10.4 long-read sequencing enables near-perfect bacterial genomes from pure cultures and metagenomes without short-read or reference polishing. BioRxiv.

