## Supplementary material for "Comparison of R9.4.1/Kit10 and R10/Kit12 Oxford Nanopore flowcells and chemistries in bacterial genome reconstruction"

**Table S1. DNA concentrations at initial extraction.** Measurements taken on 13/Sep/2021.

| **Well** | **DIN** | **Concentration (ng/microL)** | **Sample description** | **Alert** | **Observations** | **Notes** |
| --- | --- | --- | --- | --- | --- | --- |
| A1 | - | 80.0 | Ladder |  | Ladder |  |
| B1 | 9.1 | 438 | *Pseudomonas aeruginosa* PA01 | ! | Sample concentration outside functional range for DIN and the assay |  |
| C1 | 8.9 | 198 | *Klebsiella pneumoniae* MGH78578 | ! | Sample concentration outside functional range for DIN and the assay |  |
| D1 | 9.3 | 264 | Escherichia coli CFT073 | ! | Sample concentration outside functional range for DIN and the assay |  |
| E1 | 9.2 | 314 | *Staphylococcus aureus* MRSA252 - extract 1 | ! | Sample concentration outside functional range for DIN and the assay | Not used |
| F1 | 9.1 | 295 | *Staphylococcus aureus* MRSA252 - extract 2 | ! | Sample concentration outside functional range for DIN and the assay | MRSA2 was chosen for all further experiments (extracted twice due to problems with lysis) |

**Table S2. DNA concentrations at initial extraction, diluted 10x.** Measurements taken on 13/Sep/2021.

| **Well** | **DIN** | **Concentration (ng/microL)** | **Sample description** | **Alert** | **Observations** |
| --- | --- | --- | --- | --- | --- |
| A1 | - | 73.1 | Ladder |  | Ladder |
| B1 | 9.1 | 22.8 | *Pseudomonas aeruginosa* PA01 |  |  |
| C1 | 8.9 | 23.7 | *Klebsiella pneumoniae* MGH78578 |  |  |
| D1 | 9.3 | 25.0 | Escherichia coli CFT073 |  |  |
| E1 | 9.2 | 25.8 | *Staphylococcus aureus* MRSA252 - extract 1 |  | Not used |
| F1 | 9.1 | 36.4 | *Staphylococcus aureus* MRSA252 - extract 2 |  |  |

**Table S3. DNA concentrations after final experiments, taken on 31/Jan/2022.** To verify quality after long-term storage at 4°C.

| **Well** | **DIN** | **Concentration (ng/microL)** | **Sample description** | **Alert** | **Observations** | **Notes** |
| --- | --- | --- | --- | --- | --- | --- |
| A1 | - | 51.9 | Ladder |  | Ladder |  |
| B1 | 9.4 | 28.3 | *Pseudomonas aeruginosa* PA01 - diluted 10x |  |  |  |
| C1 | 9.4 | 20.8 | *Klebsiella pneumoniae* MGH78578 - diluted 10x |  |  |  |
| D1 | 9.4 | 34.1 | Escherichia coli CFT073 - diluted 10x |  |  |  |
| E1 | 9.7 | 48.9 | *Staphylococcus aureus* MRSA252 - extract 2 - diluted 10x |  |  |  |
| F1 | 9.1 | 210 | *Pseudomonas aeruginosa* Pa01 - neat DNA | ! | Sample concentration outside functional range for DIN and the assay |  |
| G1 | 9.2 | 220 | *Klebsiella pneumoniae* MGH78578 - neat DNA | ! | Sample concentration outside functional range for DIN and the assay |  |
| H1 | 9.2 | 307 | Escherichia coli CFT073 | ! | Sample concentration outside functional range for DIN and the assay |  |
| A2 | 8.7 | 426 | *Staphylococcus aureus* MRSA252 - extract 2 - neat DNA | ! | Sample concentration outside functional range for DIN and the assay |  |
| B2 | 9.1 | 238 | *Staphylococcus aureus* MRSA252 - extract 1 - neat DNA | ! | Sample concentration outside functional range for DIN and the assay | Not used |

**Table S4. Qubit DNA concentrations over time.**

|  |  | **Qubit concentration (microg/mL)** | | | |
| --- | --- | --- | --- | --- | --- |
| **Date** | **Measurement** | ***E. coli*** | ***K. pneumoniae*** | ***P. aeruginosa*** | ***S. aureus* (extract 2)** |
| 02/Sep/2021 | Post-extraction | 586 | 364 | 320 | NA |
| 03/Sep/2021 | Post-extraction | NA | NA | NA | 302 |
| 21/Sep/2021 | R10.3 sequencing | 330 | 300 | 456 | 374 |
| 23/Nov/2021 | R10.4 sequencing | 266 | 246 | NA | 296 |
| 25/Nov/2021 | R10.4 sequencing | NA | NA | 356 | NA |
| 24/Jan/2021 | Concentration check | 386 | 289 | 402 | 322 |
| 11/Feb/2022 | R10.4 multiplex sequencing | 274 | 248 | 334 | 312 |

**Table S5. Flowcell use and pore count QC for each flowcell**

| **Flow Cell number** | **Flow Cell/Chemistry** | **Pore count pre-sequencing** | **Organism** |
| --- | --- | --- | --- |
| FAR12872 | 9.4/Kit10 | 1187 | *E. coli + K. pneumoniae + P. aeruginosa + S. aureus* - multiplexed (rapid barcoding) |
| FAQ96597 | 10.3/Kit12 | 1296 | *P. aeruginosa* |
| FAQ91874 | 10.3/Kit12 | 1374 | *K. pneumoniae* |
| FAQ91081 | 10.3/Kit12 | 1250 | *E. coli* |
| FAQ96358 | 10.3/Kit12 | 1242 | *S. aureus* |
| FAR04200 | 10.4/Kit12 | 1434 | *E. coli* |
| FAR28144 | 10.4/Kit12 | 1526 | *K. pneumoniae* |
| FAR28230 | 10.4/Kit12 | 1598 | *S. aureus* |
| FAR28230 - after wash | 10.4/Kit12 | 1126 | *P. aeruginosa* |
| FAS62700 | 10.4/Kit12 | 1275 | *E. coli + K. pneumoniae + P. aeruginosa + S. aureus* - multiplexed (native barcoding) |

**Table S6. Median and modal read accuracies for all modality, flowcell/kit and basecalling combinations.** Accuracies are evaluated against the relevant Illumina-corrected reference.

| **Sequence type** | **Plexed (yes/**  **no)** | **Flowcell/**  **chemistry** | **Species** | **Read type/basecaller** | **Median % accuracy** | **Modal % accuracy** |
| --- | --- | --- | --- | --- | --- | --- |
| Illumina | Yes | NA | *E. coli* | NA | 100.0 | 100.0 |
| Nanopore | Yes | R9.4.1 | *E. coli* | hac | 96.2 | 97.2 |
| Nanopore | Yes | R9.4.1 | *E. coli* | sup | 95.8 | 96.7 |
| Nanopore | No | R10.3 | *E. coli* | hac | 96.9 | 98.1 |
| Nanopore | No | R10.3 | *E. coli* | sup | 96.4 | 97.8 |
| Nanopore | No | R10.4 | *E. coli* | hac | 95.7 | 96.5 |
| Nanopore | No | R10.4 | *E. coli* | sup | 97.5 | 98.3 |
| Nanopore | No | R10.4 | *E. coli* | duplex/sup | 99.8 | 99.9 |
| Nanopore | Yes | R10.4 | *E. coli* | hac | 97.1 | 97.8 |
| Nanopore | Yes | R10.4 | *E. coli* | sup | 98.5 | 99.0 |
| Nanopore | Yes | R10.4 | *E. coli* | duplex/sup | 99.9 | 100.0 |
| Illumina | Yes | NA | *K. pneumoniae* | NA | 100.0 | 100.0 |
| Nanopore | Yes | R9.4.1 | *K. pneumoniae* | hac | 96.2 | 97.0 |
| Nanopore | Yes | R9.4.1 | *K. pneumoniae* | sup | 95.7 | 96.7 |
| Nanopore | No | R10.3 | *K. pneumoniae* | hac | 97.4 | 98.2 |
| Nanopore | No | R10.3 | *K. pneumoniae* | sup | 97.0 | 97.9 |
| Nanopore | No | R10.4 | *K. pneumoniae* | hac | 96.1 | 97.1 |
| Nanopore | No | R10.4 | *K. pneumoniae* | sup | 97.9 | 98.5 |
| Nanopore | No | R10.4 | *K. pneumoniae* | duplex/sup | 99.9 | 99.9 |
| Nanopore | Yes | R10.4 | *K. pneumoniae* | hac | 97.3 | 97.9 |
| Nanopore | Yes | R10.4 | *K. pneumoniae* | sup | 98.6 | 99.2 |
| Nanopore | Yes | R10.4 | *K. pneumoniae* | duplex/sup | 99.9 | 99.9 |
| Illumina | Yes | NA | *P. aeruginosa* | NA | 100.0 | 100.0 |
| Nanopore | Yes | R9.4.1 | *P. aeruginosa* | hac | 96.6 | 97.2 |
| Nanopore | Yes | R9.4.1 | *P. aeruginosa* | sup | 96.5 | 97.2 |
| Nanopore | No | R10.3 | *P. aeruginosa* | hac | 98.3 | 98.9 |
| Nanopore | No | R10.3 | *P. aeruginosa* | sup | 97.9 | 98.5 |
| Nanopore | No | R10.4 | *P. aeruginosa* | hac | 96.6 | 97.4 |
| Nanopore | No | R10.4 | *P. aeruginosa* | sup | 98.4 | 98.9 |
| Nanopore | No | R10.4 | *P. aeruginosa* | duplex/sup | 99.8 | 99.9 |
| Nanopore | Yes | R10.4 | *P. aeruginosa* | hac | 97.7 | 98.2 |
| Nanopore | Yes | R10.4 | *P. aeruginosa* | sup | 99.0 | 99.4 |
| Nanopore | Yes | R10.4 | *P. aeruginosa* | duplex/sup | 99.9 | 100.0 |
| Illumina | Yes | NA | *S. aureus* | NA | 100.0 | 100.0 |
| Nanopore | Yes | R9.4.1 | *S. aureus* | hac | 97.0 | 97.8 |
| Nanopore | Yes | R9.4.1 | *S. aureus* | sup | 96.8 | 97.4 |
| Nanopore | No | R10.3 | *S. aureus* | hac | 98.3 | 99.0 |
| Nanopore | No | R10.3 | *S. aureus* | sup | 98.1 | 98.7 |
| Nanopore | No | R10.4 | *S. aureus* | hac | 97.3 | 98.0 |
| Nanopore | No | R10.4 | *S. aureus* | sup | 98.7 | 99.1 |
| Nanopore | No | R10.4 | *S. aureus* | duplex/sup | 99.9 | 100.0 |
| Nanopore | Yes | R10.4 | *S. aureus* | hac | 98.3 | 99.0 |
| Nanopore | Yes | R10.4 | *S. aureus* | sup | 99.4 | 99.9 |
| Nanopore | Yes | R10.4 | *S. aureus* | duplex/sup | 100.0 | 100.0 |

**Table S7. Number of SNPs and indels observed by sequencing data type, basecalling and assembler type.** Values representing ≤1 error/100kb of sequence are shaded in green. Numbers shown are using all read data evaluated for each modality, and for unplexed R10.4.1 runs. An average reference genome size of 4,928,392bp was used to estimate the median number of errors per 100kb of sequence.

| **Sequence type** | **Plexed (yes/**  **no)** | **Flowcell/**  **chemistry** | **Read type/basecaller** | **Assembler** | **Error type (SNPs/indels)** | **Median number of error/100kb of sequence (SNPs or indels)** |
| --- | --- | --- | --- | --- | --- | --- |
| SNPs | | | | | | |
| Illumina | Yes | NA | NA | SPAdes | SNPs | 0.71 |
| Nanopore | Yes | R9.4.1 | hac | Canu | SNPs | 1.03 |
| Nanopore | Yes | R9.4.1 | hac | Flye_hq | SNPs | 1.72 |
| Nanopore | Yes | R9.4.1 | hac | Flye_hq+medaka 1x | SNPs | 0.59 |
| Nanopore | Yes | R9.4.1 | hac | Flye_hq+medaka 2x | SNPs | 0.57 |
| Nanopore | Yes | R9.4.1 | hac | Flye_hq+medaka 3x | SNPs | 0.56 |
| Nanopore | Yes | R9.4.1 | sup | Canu | SNPs | 1.24 |
| Nanopore | Yes | R9.4.1 | sup | Flye_hq | SNPs | 2.08 |
| Nanopore | Yes | R9.4.1 | sup | Flye_hq+medaka 1x | SNPs | 0.67 |
| Nanopore | Yes | R9.4.1 | sup | Flye_hq+medaka 2x | SNPs | 0.69 |
| Nanopore | Yes | R9.4.1 | sup | Flye_hq+medaka 3x | SNPs | 0.68 |
| Nanopore+Illumina | Yes | R9.4.1 | hac | Unicycler | SNPs | 4.38 |
| Nanopore+Illumina | Yes | R9.4.1 | sup | Unicycler | SNPs | 4.38 |
| Nanopore | No | R10.3 | hac | Canu | SNPs | 0.78 |
| Nanopore | No | R10.3 | hac | Flye_hq | SNPs | 1.74 |
| Nanopore | No | R10.3 | hac | Flye_hq+medaka 1x | SNPs | 0.23 |
| Nanopore | No | R10.3 | hac | Flye_hq+medaka 2x | SNPs | 0.21 |
| Nanopore | No | R10.3 | hac | Flye_hq+medaka 3x | SNPs | 0.21 |
| Nanopore | No | R10.3 | sup | Canu | SNPs | 0.89 |
| Nanopore | No | R10.3 | sup | Flye_hq | SNPs | 1.73 |
| Nanopore | No | R10.3 | sup | Flye_hq+medaka 1x | SNPs | 0.92 |
| Nanopore | No | R10.3 | sup | Flye_hq+medaka 2x | SNPs | 0.93 |
| Nanopore | No | R10.3 | sup | Flye_hq+medaka 3x | SNPs | 0.93 |
| Nanopore | No | R10.4 | hac | Canu | SNPs | 1.71 |
| Nanopore | No | R10.4 | hac | Flye_hq | SNPs | 3.12 |
| Nanopore | No | R10.4 | hac | Flye_hq+medaka 1x | SNPs | 0.71 |
| Nanopore | No | R10.4 | hac | Flye_hq+medaka 2x | SNPs | 0.73 |
| Nanopore | No | R10.4 | hac | Flye_hq+medaka 3x | SNPs | 0.69 |
| Nanopore | No | R10.4 | sup | Canu | SNPs | 1.23 |
| Nanopore | No | R10.4 | sup | Flye_hq | SNPs | 2.77 |
| Nanopore | No | R10.4 | sup | Flye_hq+medaka 1x | SNPs | 1.02 |
| Nanopore | No | R10.4 | sup | Flye_hq+medaka 2x | SNPs | 1.06 |
| Nanopore | No | R10.4 | sup | Flye_hq+medaka 3x | SNPs | 1.02 |
| Nanopore | No | R10.4 | duplex/sup | Canu | SNPs | 0.26 |
| Nanopore | No | R10.4 | duplex/sup | Flye_hq | SNPs | 0.27 |
| Nanopore | No | R10.4 | duplex/sup | Flye_hq+medaka 1x | SNPs | 0.21 |
| Nanopore | No | R10.4 | duplex/sup | Flye_hq+medaka 2x | SNPs | 0.21 |
| Nanopore | No | R10.4 | duplex/sup | Flye_hq+medaka 3x | SNPs | 0.21 |
| Indels | | | | | | |
| Illumina | Yes | NA | NA | SPAdes | Indels | 0.02 |
| Nanopore | Yes | R9.4.1 | hac | Canu | Indels | 26.83 |
| Nanopore | Yes | R9.4.1 | hac | Flye_hq | Indels | 17.57 |
| Nanopore | Yes | R9.4.1 | hac | Flye_hq+medaka 1x | Indels | 2.97 |
| Nanopore | Yes | R9.4.1 | hac | Flye_hq+medaka 2x | Indels | 3.37 |
| Nanopore | Yes | R9.4.1 | hac | Flye_hq+medaka 3x | Indels | 3.00 |
| Nanopore | Yes | R9.4.1 | sup | Canu | Indels | 25.83 |
| Nanopore | Yes | R9.4.1 | sup | Flye_hq | Indels | 17.56 |
| Nanopore | Yes | R9.4.1 | sup | Flye_hq+medaka 1x | Indels | 3.17 |
| Nanopore | Yes | R9.4.1 | sup | Flye_hq+medaka 2x | Indels | 3.23 |
| Nanopore | Yes | R9.4.1 | sup | Flye_hq+medaka 3x | Indels | 3.13 |
| Nanopore+Illumina | Yes | R9.4.1 | hac | Unicycler | Indels | 0.56 |
| Nanopore+Illumina | Yes | R9.4.1 | sup | Unicycler | Indels | 0.57 |
| Nanopore | No | R10.3 | hac | Canu | Indels | 3.22 |
| Nanopore | No | R10.3 | hac | Flye_hq | Indels | 1.79 |
| Nanopore | No | R10.3 | hac | Flye_hq+medaka 1x | Indels | 0.92 |
| Nanopore | No | R10.3 | hac | Flye_hq+medaka 2x | Indels | 0.50 |
| Nanopore | No | R10.3 | hac | Flye_hq+medaka 3x | Indels | 0.44 |
| Nanopore | No | R10.3 | sup | Canu | Indels | 5.90 |
| Nanopore | No | R10.3 | sup | Flye_hq | Indels | 3.01 |
| Nanopore | No | R10.3 | sup | Flye_hq+medaka 1x | Indels | 1.17 |
| Nanopore | No | R10.3 | sup | Flye_hq+medaka 2x | Indels | 1.13 |
| Nanopore | No | R10.3 | sup | Flye_hq+medaka 3x | Indels | 1.10 |
| Nanopore | No | R10.4 | hac | Canu | Indels | 9.94 |
| Nanopore | No | R10.4 | hac | Flye_hq | Indels | 4.57 |
| Nanopore | No | R10.4 | hac | Flye_hq+medaka 1x | Indels | 1.00 |
| Nanopore | No | R10.4 | hac | Flye_hq+medaka 2x | Indels | 0.98 |
| Nanopore | No | R10.4 | hac | Flye_hq+medaka 3x | Indels | 0.98 |
| Nanopore | No | R10.4 | sup | Canu | Indels | 1.54 |
| Nanopore | No | R10.4 | sup | Flye_hq | Indels | 0.99 |
| Nanopore | No | R10.4 | sup | Flye_hq+medaka 1x | Indels | 0.41 |
| Nanopore | No | R10.4 | sup | Flye_hq+medaka 2x | Indels | 0.41 |
| Nanopore | No | R10.4 | sup | Flye_hq+medaka 3x | Indels | 0.41 |
| Nanopore | No | R10.4 | duplex/sup | Canu | Indels | 0.37 |
| Nanopore | No | R10.4 | duplex/sup | Flye_hq | Indels | 0.48 |
| Nanopore | No | R10.4 | duplex/sup | Flye_hq+medaka 1x | Indels | 0.18 |
| Nanopore | No | R10.4 | duplex/sup | Flye_hq+medaka 2x | Indels | 0.18 |
| Nanopore | No | R10.4 | duplex/sup | Flye_hq+medaka 3x | Indels | 0.18 |

**Figure S1. DNA extract tapestation profile.** Neat DNA, measurements taken on 13/Sep/2021.

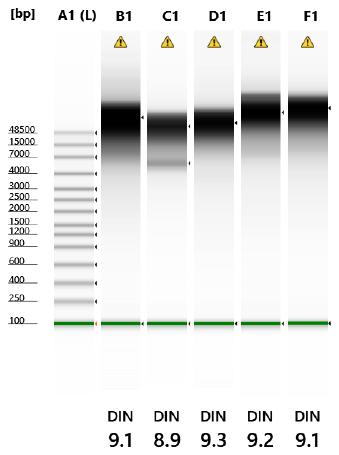

**Figure S2. DNA extract tapestation profile.** DNA diluted 10x, measurements taken on 13/Sep/2021.

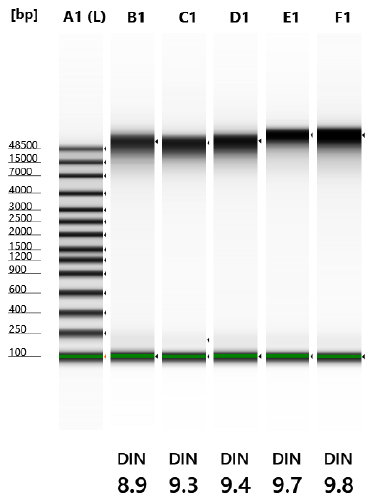

**Figure S3. DNA extract tapestation profile.** DNA diluted 10x, measurements taken on 31/Jan/2022.

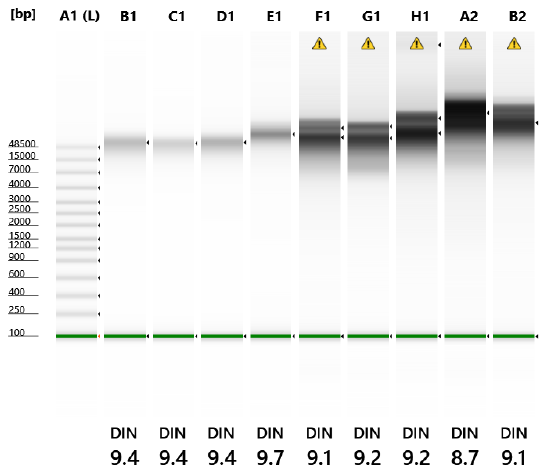

**Figure S4. Sequencing yields over time for unplexed/multi(plexed) runs stratified by species and for R10.4, duplex versus simplex read outputs.** Note y-axis scales (depicting yields in cumulative Gb) are different across facets.

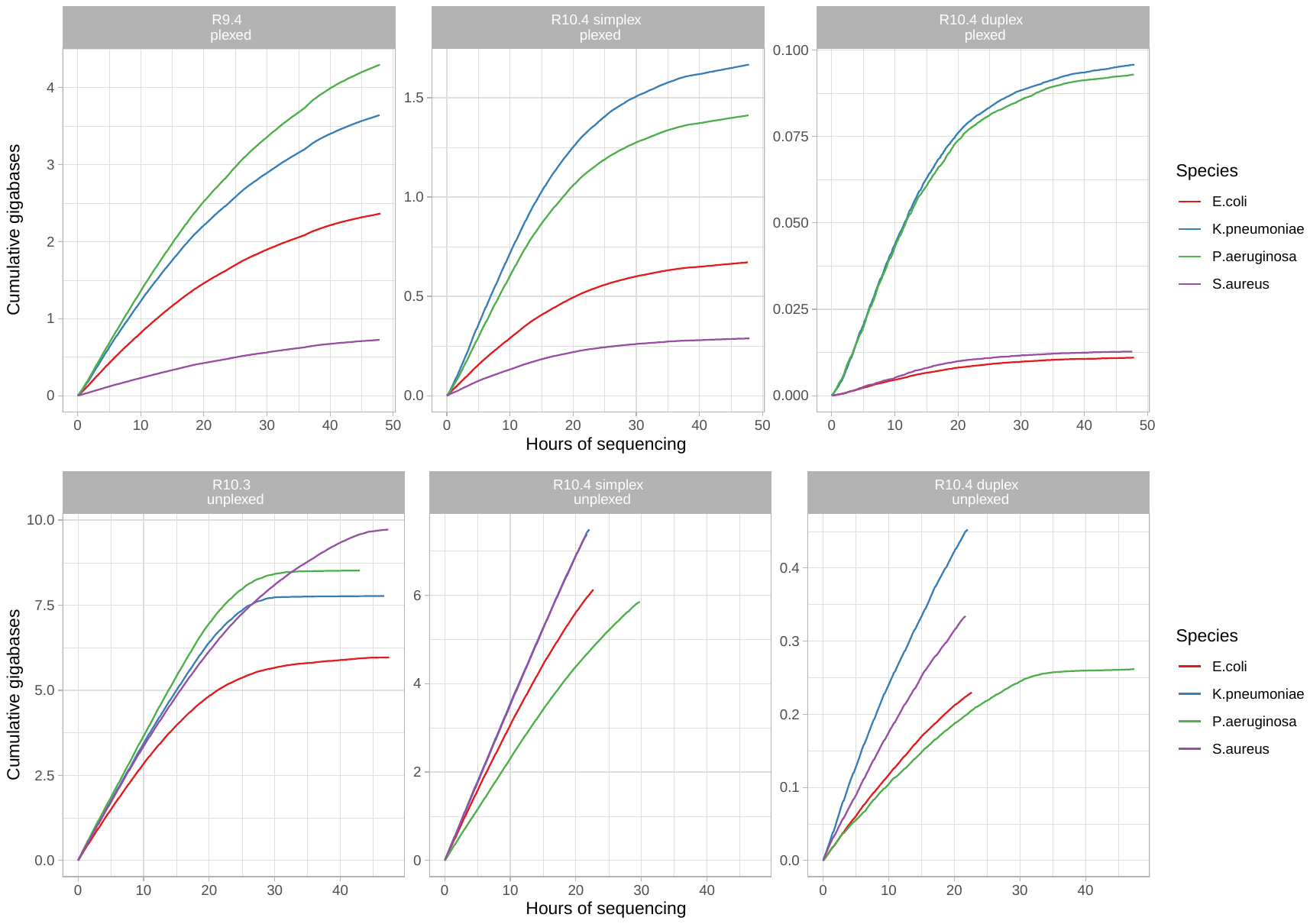

**Figure S5. Impact of sub-sampling on number of contigs assembled for each species, modality and assembler - (A) chromosomes and (B) *K. pneumoniae* plasmids.** In the plasmid plots, plasmids are denoted by their length in bp in the facet headers. For each comparison, there should be seven sub-sampling levels (represented by the rainbow colours) - if any of these are absent, then that contig was missing in the assembly. Similarly, in all plots, there should be single unique contig assembled (i.e. the y-axis should be 1). If this is >1, then the assembly is fragmented; or if 0, then the contig is absent.

**A.**

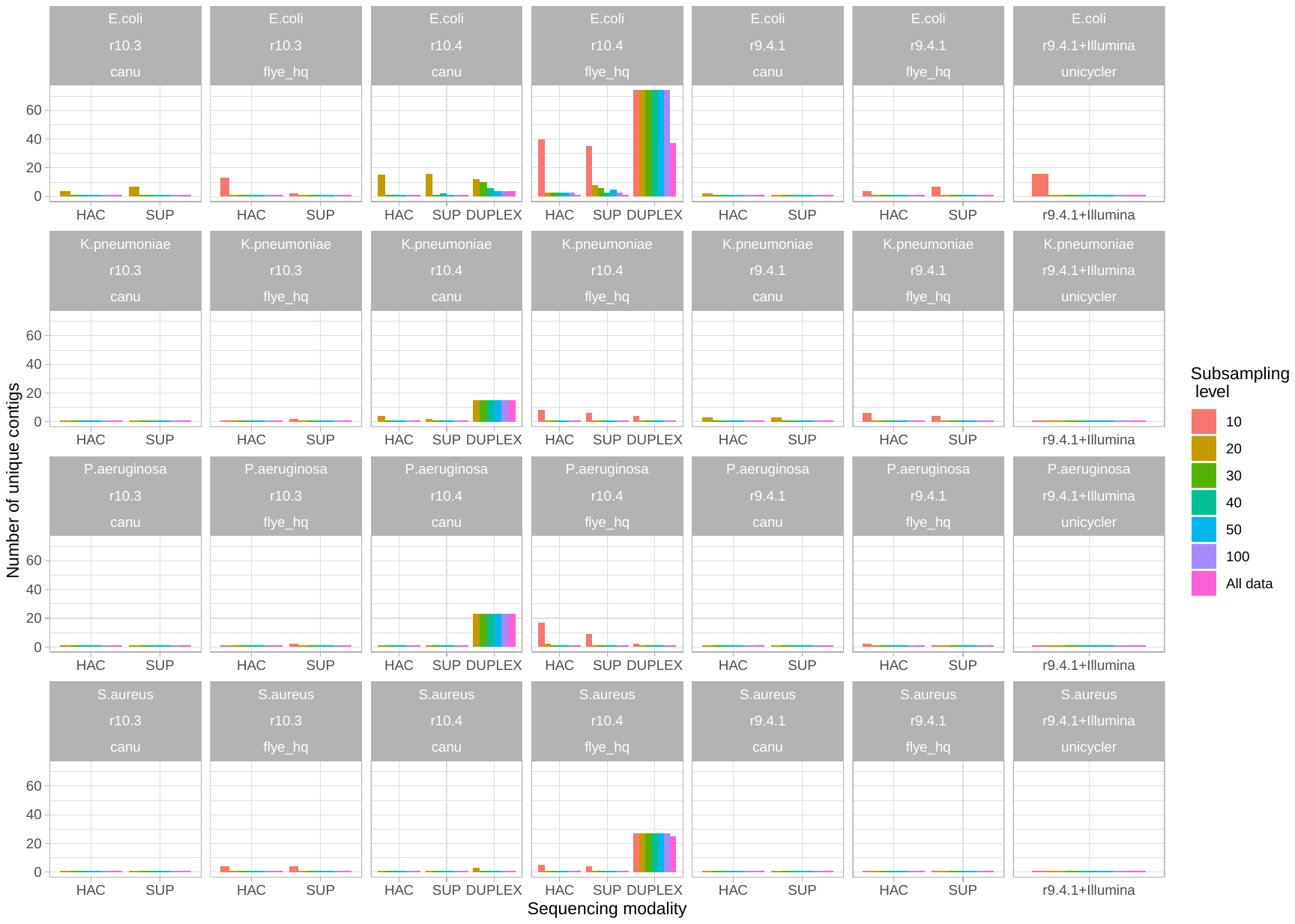

**B.**

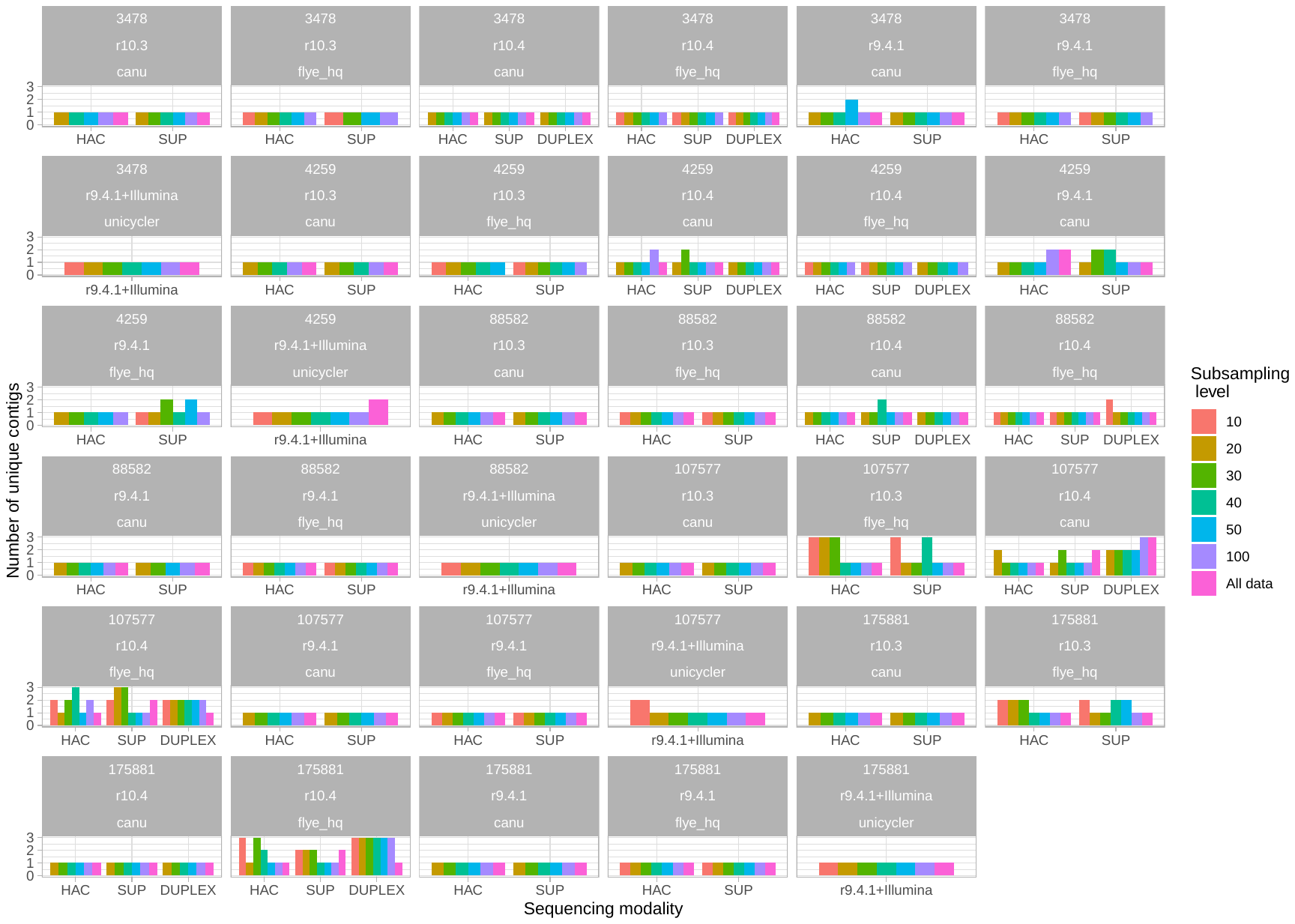

**Figure S6. Impact of Medaka polishing of Flye assemblies, by species and read sub-sampling stratum.**

**A. *E. coli***

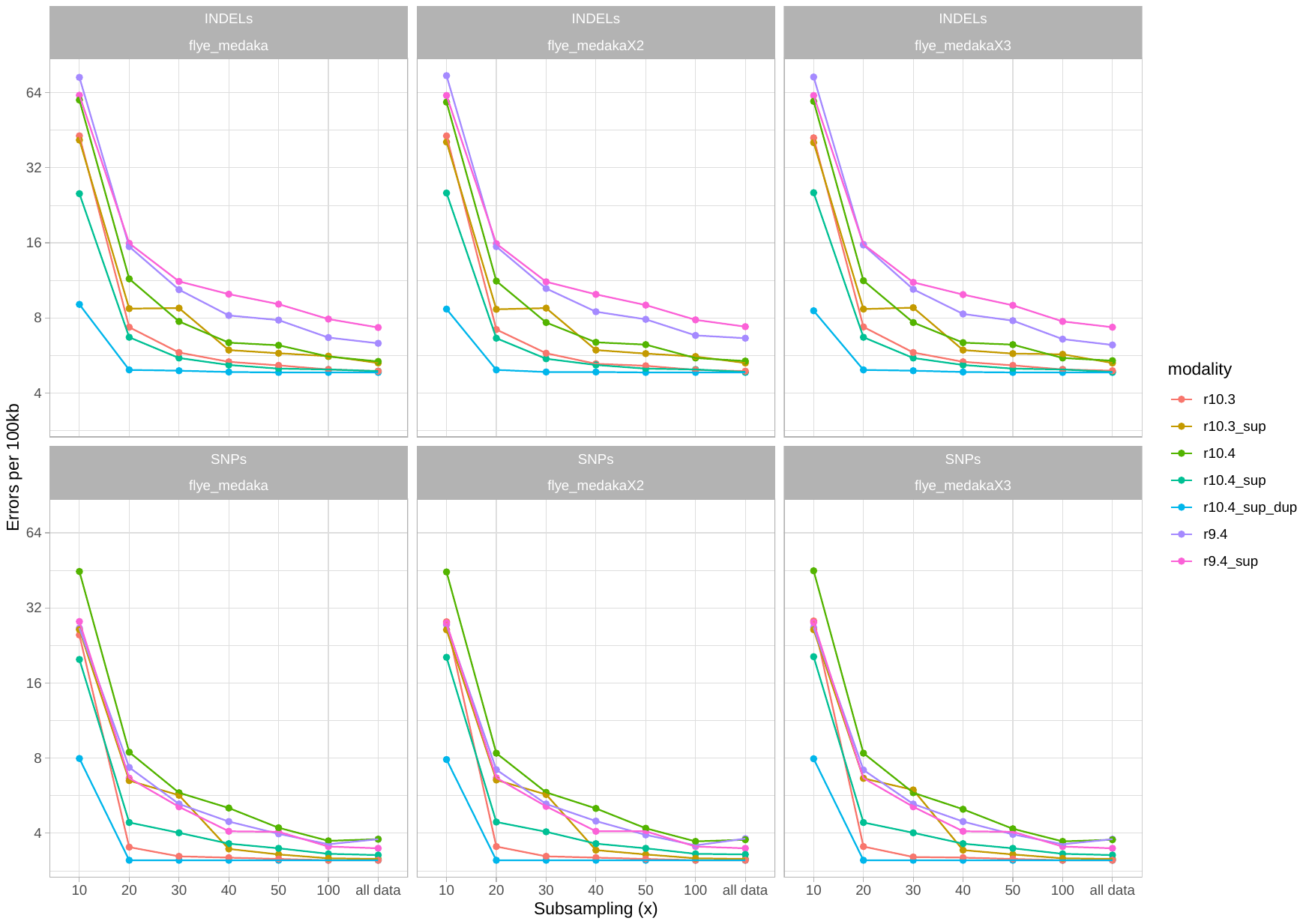

**B. *K. pneumoniae***

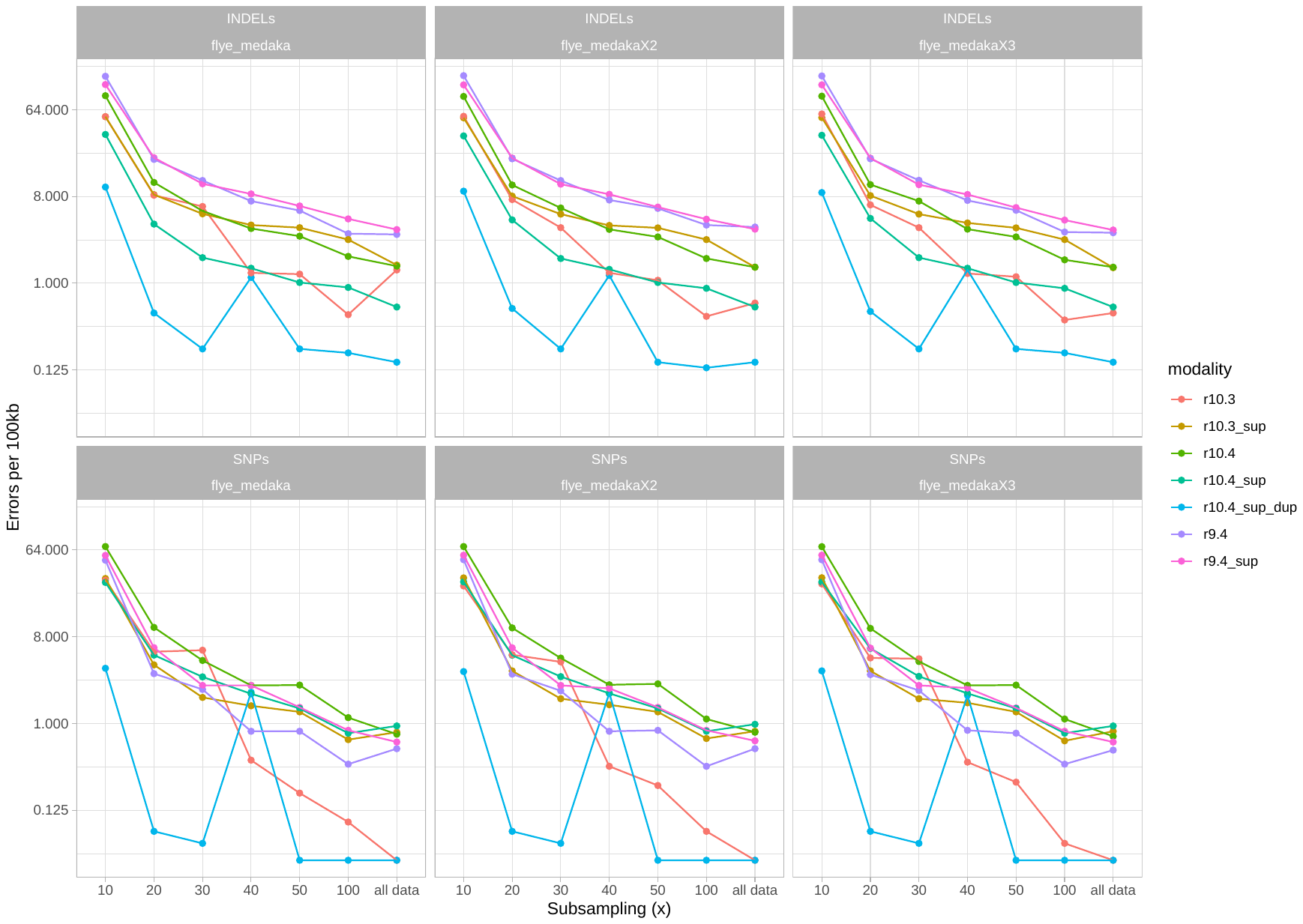

**C. *P. aeruginosa***

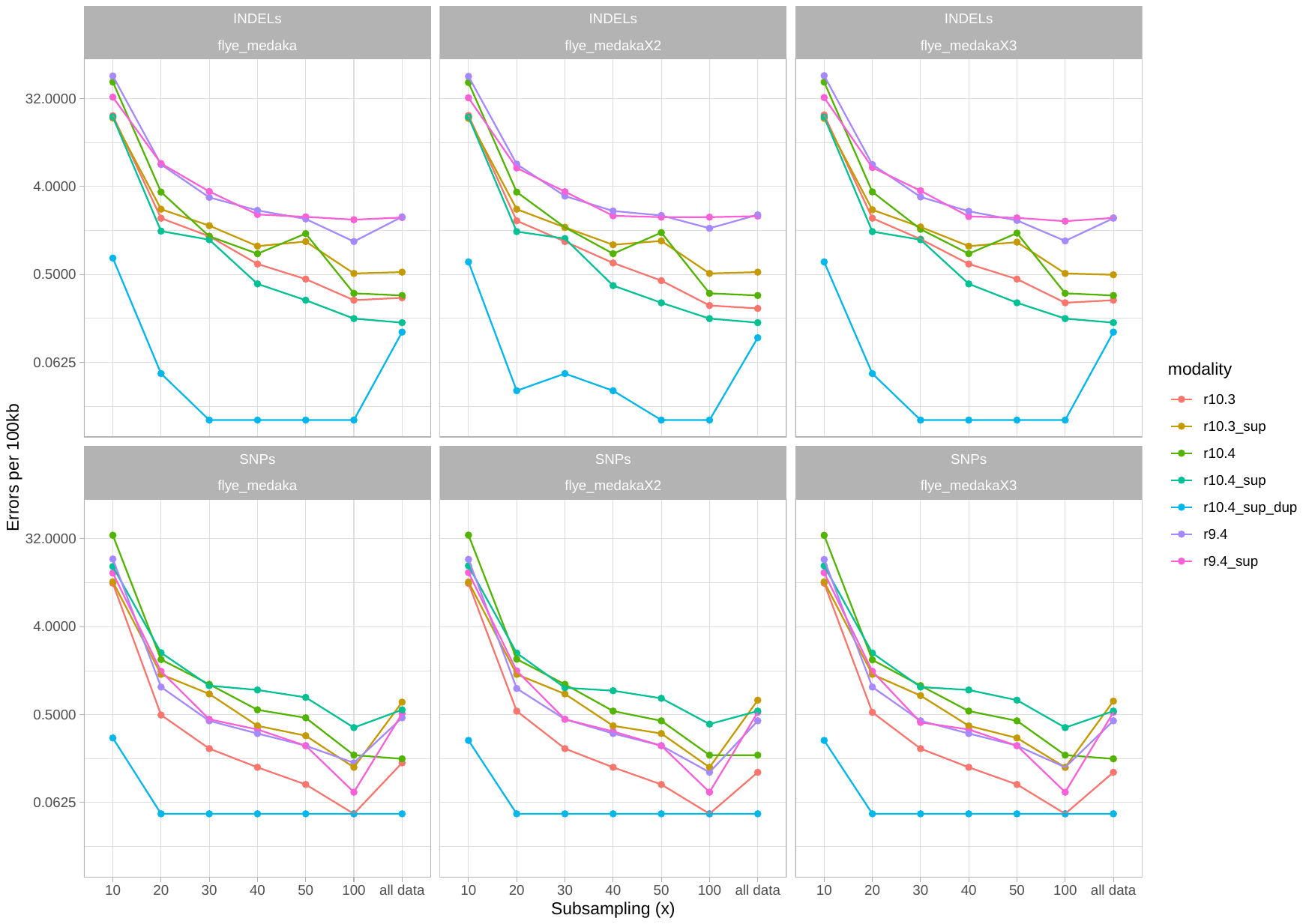

**D. *S. aureus***

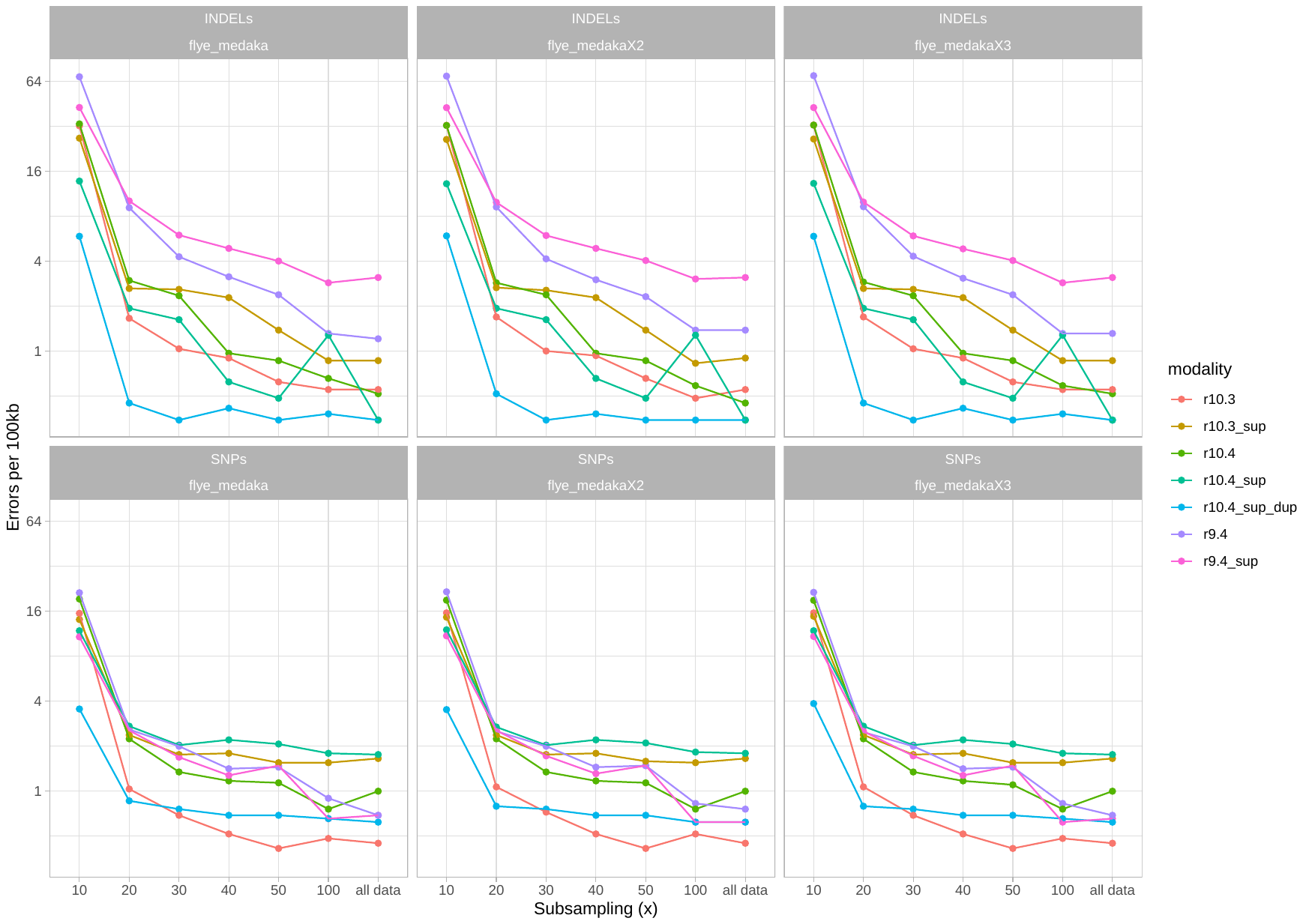
